# Botulinum neurotoxin accurately separates tonic vs phasic transmission and reveals heterosynaptic plasticity rules in Drosophila

**DOI:** 10.1101/2022.02.15.480429

**Authors:** Yifu Han, Chun Chien, Pragya Goel, Kaikai He, Cristian Pinales, Chris Buser, Dion Dickman

## Abstract

In developing and mature nervous systems, diverse neuronal subtypes innervate common targets to establish and maintain functional neural circuits. A major challenge towards understanding the structural and functional architecture of neural circuits is to separate these inputs and determine their intrinsic and heterosynaptic relationships. The *Drosophila* larval neuromuscular junction is a powerful model system to study these questions, where two glutamatergic motor neurons, the strong phasic-like **Is** and weak tonic-like **Ib**, co-innervate individual muscle targets to coordinate locomotor behavior. However, complete neurotransmission from each input has never been electrophysiologically separated. We have developed a botulinum neurotoxin, BoNT-C, that eliminates both spontaneous and evoked neurotransmission without perturbing synaptic growth or structure, enabling the first approach that accurately isolates input-specific neurotransmission. Selective expression of BoNT-C in Is or Ib motor inputs disambiguates functional properties of each input. Importantly, the composite values of Is and Ib neurotransmission can be fully recapitulated by isolated physiology from each input. Finally, selective silencing by BoNT-C does not induce heterosynaptic structural or functional plasticity at the convergent input. Thus, BoNT-C establishes the first approach to cleanly separate neurotransmission between tonic vs phasic neurons and defines heterosynaptic plasticity rules in a powerful model glutamatergic circuit.

## INTRODUCTION

Neural circuits are established in development, modified by experience, and must be maintained at stable physiologic levels throughout the lifetime of an organism. Most cells embedded in neural circuits are innervated by multiple neurons, which differ in the number and strength of synaptic connections, the classes of neurotransmitters and neuropeptides released, and their patterns of activity. There is also evidence that changes in synapse number or function at one input can induce adaptive or Hebbian modulations in transmission at convergent inputs, termed heterosynaptic plasticity (Aponte-Santiago et al., 2020; Chater and Goda, 2021; Dittman and Regehr, 1997; Wang et al., 2021). The *Drosophila* larval neuromuscular junction (NMJ) is a powerful model system for revealing fundamental principles of transmission and heterosynaptic plasticity, where synaptic development, growth, function, and plasticity has been studied for over 40 years (Brunner and O’Kane, 1997; Charng et al., 2014; Harris and Littleton, 2015). Most muscles at this glutamatergic synapse are co-innervated by two distinct motor neurons (MNs) that coordinate muscle contraction to drive locomotor behavior, termed MN-Is and MN-Ib. These MNs differ in both structural and functional properties, with the strong MN-Is firing with phasic-like patterns and depressing with repeated stimulation, while the weak MN-Ib fires with tonic-like patterns and facilitates (Aponte-Santiago and Littleton, 2020; Hoang and Chiba, 2001; Lnenicka and Keshishian, 2000; Lu et al., 2016; Newman et al., 2017). The vast majority of fly NMJ studies of synaptic structure have focused on MN-Ib terminals, given their large relative size and amenability to light and super resolution imaging (Lu and Lichtman, 2007; Maglione and Sigrist, 2013; Sigrist and Sabatini, 2012). Similarly, most studies using electrophysiology in this system have recorded an ambiguous blend of miniature events originating from both MN-Is and MN-Ib, as well as a composite evoked response from simultaneous stimulation of both inputs. It bears emphasizing that this compound evoked response does not reflect the physiology of any actual existent synapse. This failure to fully disambiguate transmission from the strong MN-Is and weak MN-Ib has limited our understanding and ability to interpret synaptic transmission and plasticity at the *Drosophila* NMJ.

Although most electrophysiological studies using the *Drosophila* NMJ have failed to cleanly separate transmission from entire MN-Is and MN-Ib inputs, important insights have been gleaned into their respective properties. Early anatomical studies characterized their relative size and structures, which established that MN-Is boutons are smaller and contain fewer release sites, while those of MN-Ib are larger and opposed by a more elaborate subsynaptic reticulum (Atwood et al., 1993; Jia et al., 1993; Lnenicka and Keshishian, 2000; Schuster et al., 1996). Differences were also observed in the relative abundance of postsynaptic glutamate receptor subtypes at MN-Is vs MN-Ib NMJs (Marrus et al., 2004; Schmid et al., 2008). Macro patch recordings at identified boutons suggested spontaneous events were larger at MN-Is terminals (Pawlu et al., 2004), consistent with electron microscopy studies that showed synaptic vesicles were larger at MN-Is terminals (Karunanithi et al., 2002). Finally, threshold stimulus manipulations established that single action-potential evoked transmitter release from MN-Is drives most of the depolarization in the postsynaptic muscle (Lu et al., 2016). Together, this important body of work detailed fundamental properties of the two MN subtypes in *Drosophila*.

Recent innovations in Ca^2+^ imaging and the identification of GAL4 drivers that selectively express in MN-Is vs MN-Ib has enabled new attempts to isolate transmission between the two inputs. First, postsynaptic Ca^2+^ sensors were developed that revealed important differences in quantal release events, active zone-specific release characteristics, and plasticity between MN-Is vs MN-Ib NMJs (Akbergenova et al., 2018; Gratz et al., 2019; Newman et al., 2022, 2017; Peled and Isacoff, 2011). Following the recent identification of MN-Is and MN-Ib GAL4 drivers (Aponte-Santiago et al., 2020; Pérez-Moreno and O’Kane, 2019), input-specific genetic ablation suggested heterosynaptic structural plasticity could be induced between convergent Is vs Ib inputs (Aponte-Santiago et al., 2020; Wang et al., 2021). However, to what extent heterosynaptic functional plasticity was expressed was unclear due to an inability to isolate transmission from either input. Finally, selective optogenetic stimulation of MN-Is or MN-Ib has been used to estimate evoked neurotransmission (Genç and Davis, 2019; Sauvola et al., 2021), but is unable to resolve input-specific miniature transmission and may suffer from confounds due to chronic channel rhodopsin expression in MNs. An accurate electrophysiological understanding of transmission from entire MN-Is vs MN-Ib NMJs has therefore remained unresolved due to the significant limitations these approaches. Similarly, clear rules and mechanistic insights into the signaling systems mediating heterosynaptic plasticity at the fly NMJ has not been established.

We have therefore sought to develop an optimal approach to cleanly separate neurotransmission between tonic and phasic motor inputs in *Drosophila*. Towards this goal, we screened a series of botulinum toxins for abilities to prevent neurotransmitter release from MNs. This approach identified Botulinum NeuroToxin-C (BoNT-C) to eliminate all spontaneous and evoked neurotransmitter release without imposing apparent intrinsic toxicity or plasticity at the convergent input. BoNT-C now enables the accurate separation of motor inputs and establishes heterosynaptic plasticity rules at the *Drosophila* NMJ.

## RESULTS

### Suboptimal approaches to disambiguate tonic and phasic inputs at the *Drosophila* NMJ

An ideal approach to functionally isolate MN-Is and MN-Ib would silence all neurotransmission from one input without inducing structural or functional plasticity from the convergent input (“heterosynaptic plasticity”). If such an approach were successful, then the frequency of miniature transmission should be reduced when either input is silenced compared to recordings from wild-type NMJs (where miniature transmission is blended), and quantal size should be increased at MN-Is NMJs relative to MN-Ib. In addition, synaptic strength, as assessed by single action-potential stimulation, should be increased when evoked from MN-Is relative to MN-Ib NMJs. Importantly, when the composite values of miniature and evoked transmission from MN-Is and MN-Ib NMJs are averaged or summed, they should fully recapitulate all aspects of neurotransmission as assessed from standard wild-type NMJ recordings, where miniature events are a mix from both inputs and evoked transmission is an ambiguous average of simultaneous release from both MN-Is and MN-Ib.

The fly larval musculature is composed of 30 repeated muscle segments innervated by 36 distinct MNs: ∼30 MN-Ib and 3 MN-Is (Arzan Zarin and Labrador, 2017; Clark et al., 2018; Hoang and Chiba, 2001; Kim et al., 2009; Lnenicka and Keshishian, 2000). MN-Ib inputs typically innervate single muscles, while MN-Is co-innervates groups of several muscles (Fig. S1). To distinguish transmission between MN-Is and MN-Ib inputs at the *Drosophila* NMJ, we first confirmed expression profiles of four MN GAL4 drivers that express at the muscle 6/7 NMJ, the primary NMJ that has been used for electrophysiology in the field and that we will focus on in this study: 1) *OK6-GAL4*, which drives GAL4 expression in all MNs, including the ones that innervate muscle 6/7; 2) *OK319-GAL4*, which expresses in small subsets of both Is and Ib MNs, including the ones that innervate muscle 6/7; 3) “Is-GAL4” (*dHB9-GAL4*), which expresses in the MN-Is that innervates muscle 6/7; and 4) “Ib-GAL4” (*R27E09-GAL4*), which expresses in the single MN-Ib that innervates muscle 6/7. Expression of these drivers proved specific (Fig. S1), motivating a genetic dissection of the MN-Ib and MN-Is inputs at the muscle 6/7 NMJ.

One promising approach to electrophysiologically separate MN-Is and MN-Ib would be to employ a conditional null allele of the *vesicular glutamate transporter* (*vGlut*) that was recently developed (Banerjee et al., 2021; Sherer et al., 2020). In principle, this would be an ideal approach because all glutamate release would be silenced without otherwise perturbing innervation or synaptogenesis. Indeed, conditional loss of *vGlut* eliminates all neurotransmission in adult NMJs (Banerjee et al., 2021). However, at larval NMJs we found vGlut expression to be only moderately reduced at presynaptic terminals and did not eliminate transmission, despite trying all four of the GAL4 drivers described above to conditionally remove *vGlut* (Fig. S2). This is likely due to perdurance of maternally contributed *vGlut* and early *vGlut* synthesis, as well as the finding that an individual vGlut transporter is sufficient to fill a synaptic vesicle (Daniels et al., 2006). Hence, conditional loss of *vGlut* in larval MNs was not an effective approach for our goal of eliminating miniature and evoked transmission.

Using the Is-GAL4 and Ib-GAL4 drivers described above, we tried three alternative approaches to distinguish transmission between these two inputs. Each of these were previously used in attempts to functionally separate MN-Is/MN-Ib transmission, and each proved inadequate for our purposes. First, we used selective optogenetic stimulation of MN-Ib or MN-Is as recently employed (Genç and Davis, 2019; Sauvola et al., 2021). From the outset, we knew the limitations of this approach in failing to distinguish input-specific miniature transmission and an inability to perform repeated trains of stimulation to assess short-term plasticity and to resolve vesicle pools. We expressed *channel rhodopsin* (ChR) in all MNs (OK6>ChR) or selectively in either MN-Is or MN-Ib (Fig. 1A,B), as performed in these previous studies. As expected, miniature frequency and amplitude were indistinguishable in Is>ChR and Ib>ChR compared to OK6>ChR or wild type (Fig. 1A,B), while evoked release was stronger at Is compared to Ib, as expected. However, we also noted significant differences between optogenetic vs electrical stimulation (Fig. 1B,D; Fig. S3), where baseline evoked EPSP values was reduced by ∼33% OK6>ChR compared to wild type controls, suggesting chronic expression of ChR alone induces significant changes in intrinsic excitability and synaptic function. Next, we expressed tetanus toxin (TNT; (Sweeney et al., 1995)) in an attempt to selectively silence evoked transmission in either MN (Fig. 1C,D). Although TNT expression blocks evoked release, miniature transmission persists (Choi et al., 2014; Sweeney et al., 1995), so we also knew from the outset that TNT, like optogenetic stimulation, would fail to separate miniature transmission. Indeed, selective expression of TNT does not significantly change mEPSP frequency or amplitude from wild type (Fig. 1C,D). Further, while evoked EPSPs were larger at MN-Is NMJs compared to MN-Ib, as observed with optogenetic stimulation, input-specific TNT expression was reported to induce heterosynaptic plasticity (Aponte-Santiago et al., 2020), making it unclear if the input-specific differences in synaptic strength reflects genuine wild-type behavior. Finally, we tried genetic ablation of either MN-Is or MN-Ib by expression of the pro-apoptotic genes *reaper* (*rpr*) and *head involution defective* (*hid*). Is>rpr.hid cleanly eliminated MN-Is inputs (Fig. S4), but was also reported to induce heterosynaptic plasticity (Aponte-Santiago et al., 2020; Wang et al., 2021). However, Ib>rpr.hid did not always completely ablate MN-Ib inputs, yet always fully eliminated all MN-Is inputs (Fig. S4). While electrophysiological recordings from Is>rpr.hid appeared to silence transmission from MN-Ib (Fig. 1E,F), virtually all transmission was ablated at Ib>rpr.hid NMJs (Fig. 1E,F). Thus, conditional mutagenesis, optogenetics, TNT expression, and genetic ablation each proved inadequate to accurately distinguish input-specific transmission.

**Figure 1:**
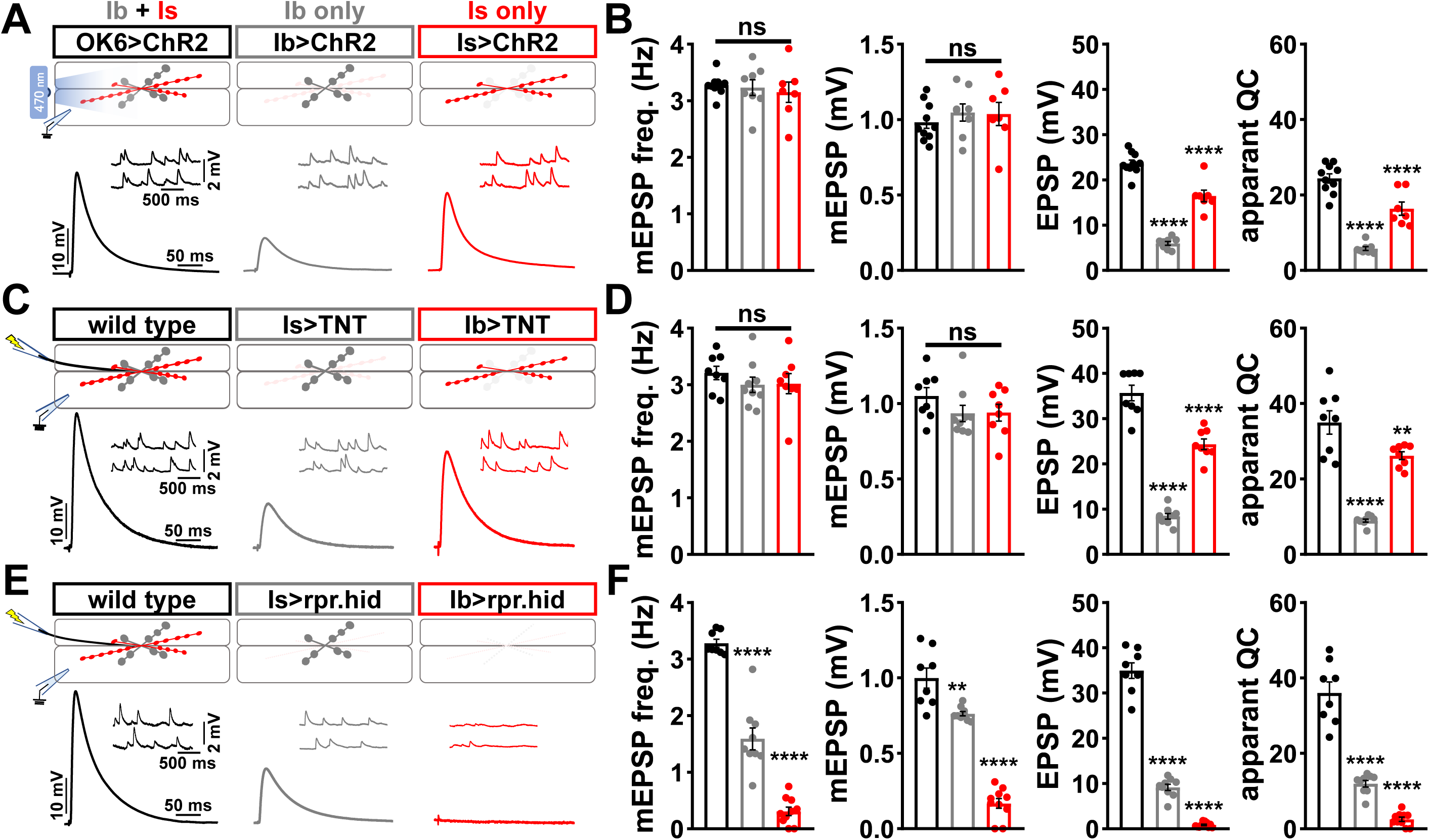
Suboptimal approaches for selectively isolating tonic and phasic neurotransmission at the *Drosophila* NMJ. **(A)** Schematic of recording configuration and representative mEPSP and EPSP traces illustrating that overexpression of *ChR2^T159C^* in MN-Ib or -Is enables optically evoked EPSP events from either MN-Ib or -Is NMJs. Genotypes: OK6>ChR2 (*w*;*OK6-GAL4*/*UAS-ChR2^T159C^*;*+*); Ib>ChR2 (*w*;*UAS-ChR2^T159C^*/*+*;*dHB9-GAL4*/*+*); Is>ChR2 (*w*;*UAS-ChR2^T159C^*/*+*;*R27E09-GAL4*/*+*). **(B)** Quantification of mEPSP frequency, amplitude, EPSP, and apparent quantal content values in the indicated genotypes in (A). Note that because input-specific mEPSP values cannot be determined using this optogenetic approach, inaccurate quantal content values are shown by simply dividing by the EPSP values by the same averaged mEPSP values. **(C)** Schematic and representative traces following selective expression of tetanus toxin (TNT) in MN-Ib or -Is. Genotypes: wild type (*w^1118^*); Is>TNT (*w*;*+*;*R27E09-GAL4*/*UAS-TNT*); Ib>TNT (*w*;*+*;*dHB9-GAL4*/*UAS-TNT*). **(D)** Quantification of the indicated values of the genotypes shown in (C), with the same inaccuracies in determining quantal content. **(E)** Schematic and representative traces following input-specific expression of the pro-apoptotic genes *reaper* (*rpr*) and *head-involution-defective* (*hid*). Note that MN-Is NMJs are completely absent following MN-Ib ablation. Genotypes: wild type (*w*); Is>rpr.hid (*UAS-rpr.hid*/*+*;*+*; *R27E09-GAL4*/+); Ib>rpr.hid (*UAS-rpr.hid*/*+*;*+*; *dHB9-GAL4*/*+*). **(F)** Quantification of the indicated values of the genotypes shown in (E). Error bars indicate ±SEM. ****P<0.0001; ***P<0.001; **P<0.01; ns, not significant. Additional statistical details are shown in Table S2.

### BoNT-C eliminates neurotransmission without confounding alterations in pre- or post- synaptic structure

We therefore sought to develop a new approach designed to eliminate all neurotransmission (both miniature and evoked release) and that ideally would not otherwise perturb MN structure or innervation. This search led us to Botulinum NeuroToxins (BoNTs), clostridial toxins that function as potent protein enzymes to cleave the synaptic vesicle SNARE complexes necessary for exocytosis (Fig. 2A,B; (Dong et al., 2019)). We cloned sequences encoding four different BoNT light chains (BoNT-A, -B, -C, and -E), each targeting distinct SNARE components (Fig. 2A,B) into *Drosophila* transgenic vectors under control of GAL4-responsive UAS sequences. Importantly, the cleavage targets of each BoNT were validated for the *Drosophila* proteins (Backhaus et al., 2016). From the outset, we suspected that BoNT-C would likely be an ideal candidate, since it targets the t-SNARE Syntaxin, mutations of which in *Drosophila* eliminate all miniature and evoked release (Deitcher et al., 1998; Schulze et al., 1995). As a basis of comparison, miniature transmission persists mutations in the *Drosophila* vSNARE *neuronal synaptobrevin* (*n-syb;* (Broadie et al., 1995; Deitcher et al., 1998)), and n-Syb is the target of TNT (n-Syb; Fig. 2A). As an initial screen for SNARE cleavage activity, we evaluated lethality following pan-neuronal (*C155-GAL4*) and MN-specific (*OK6-GAL4*) BoNT expression. Following these filters, only BoNT-B and BoNT-C caused embryonic lethality (Fig. 2C). To circumvent lethality, we then expressed either BoNT-B and BoNT-C with *OK319-GAL4*, where NMJ electrophysiological recordings revealed BoNT-B expression failed to fully eliminate transmission (Fig. 2C, Table S1). However, while miniature transmission persists in OK319>TNT, OK319>BoNT-C eliminated both spontaneous and evoked neurotransmission (Fig. 2D,E). Thus, BoNT-C expression in MNs was identified as the only SNARE toxin capable of eliminating both miniature and evoked neurotransmission.

**Figure 2:**
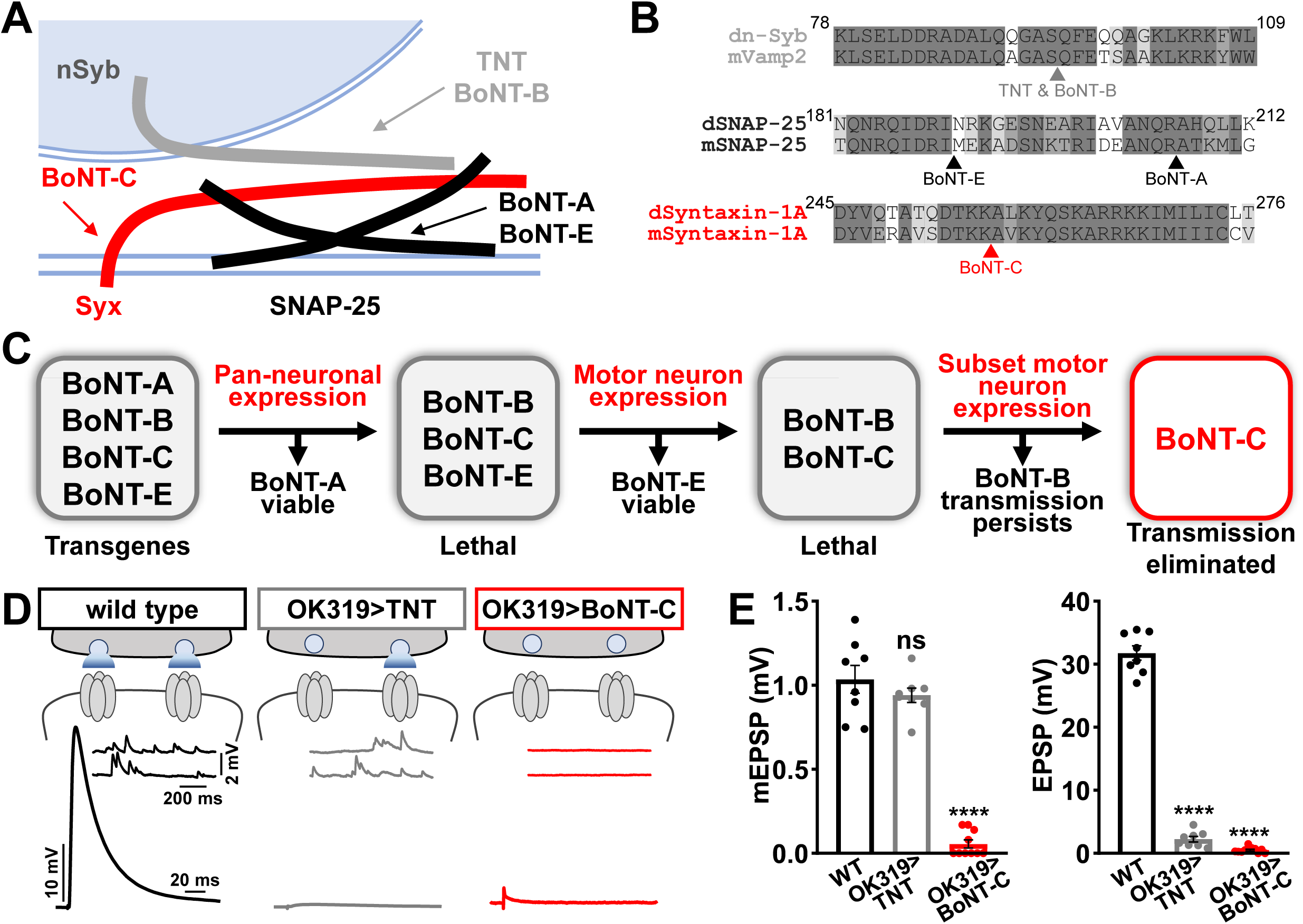
BoNT-C eliminates both spontaneous and evoked transmission. **(A)** Schematic of synaptic SNARE proteins that are targets for enzymatic cleavage by the Tetanus (TNT) and Botulinum (BoNT) neurotoxins. **(B)** Amino acid sequence alignments of the cleavage sites for the indicated TNT and BoNT toxins in the mouse and Drosophila SNARE components Synaptobrevin (Vamp2 and Neuronal Synaptobrevin (dN-Syb)), SNAP25, and Syntaxin (Syntaxin-1A). Arrows illustrate the conserved cleavage sites of TNT, BoNT-A, BoNT-B, BoNT-C, and BoNT-E. **(C)** BoNT screening flow chart: Four BoNT lines were first tested for lethality when expressed using the pan-neuronal *c155-GAL4* driver, then similarly tested when crossed to the motor neuron-specific *OK6-GAL4* driver, resulting in only BoNT-B and BoNT-C causing lethality. These transgenes were then crossed to the *OK319-GAL4* driver, which expressed in only a subset of motor neurons, and electrophysiological recordings revealed that while transmission persisted in BoNT-B, transmission was completed blocked in BoNT-C. **(D)** Schematic and representative electrophysiological traces illustrating that while miniature transmission persists at NMJs poisoned by TNT expression, transmission was completely blocked at NMJs expressing BoNT-C. **(E)** Quantification of mEPSP and EPSP amplitudes in the indicated genotypes: wild type (*w*); OK319>TNT (*w*;*OK319-GAL4*/*UAS-TNT*;+); OK319>BoNT-C (*w*;*OK319-GAL4*/*+*;*UAS-BoNT-C*/+). Error bars indicate ±SEM. ****P<0.0001; ns, not significant. Additional BoNT screening results and statistical details are shown in Tables S1 and S2.

Next, we examined whether BoNT-C expression perturbs presynaptic growth or structure when expressed in MNs. Neuronal toxicity has been reported for some classes of neurotoxins (Peng et al., 2013), and we continued to use TNT expression as a comparison. Expression of TNT in both MN-Is and -Ib by OK319>GAL4 led to a ∼20% reduction in bouton and active zone number at terminals of MN-Ib (indicated by immunostaining of the active zone scaffold Bruchpilot (BRP)), while and inverse change in synaptic growth was observed in MN-Is (Fig. 3A), consistent with previous reports (Goel and Dickman, 2018; Li et al., 2018). In contrast, BoNT-C expression in both MN-Is and MN-Ib did not significantly change bouton or BRP puncta number, nor was BRP intensity altered in either input (Fig. 3B). Similarly, we observed no differences in NMJ ultrastructure following BoNT-C expression, with no significant changes in T-bar or active zone length or synaptic vesicle density (Fig. 3C,D). Thus, BoNT-C expression does not perturb presynaptic growth or structure.

**Figure 3:**
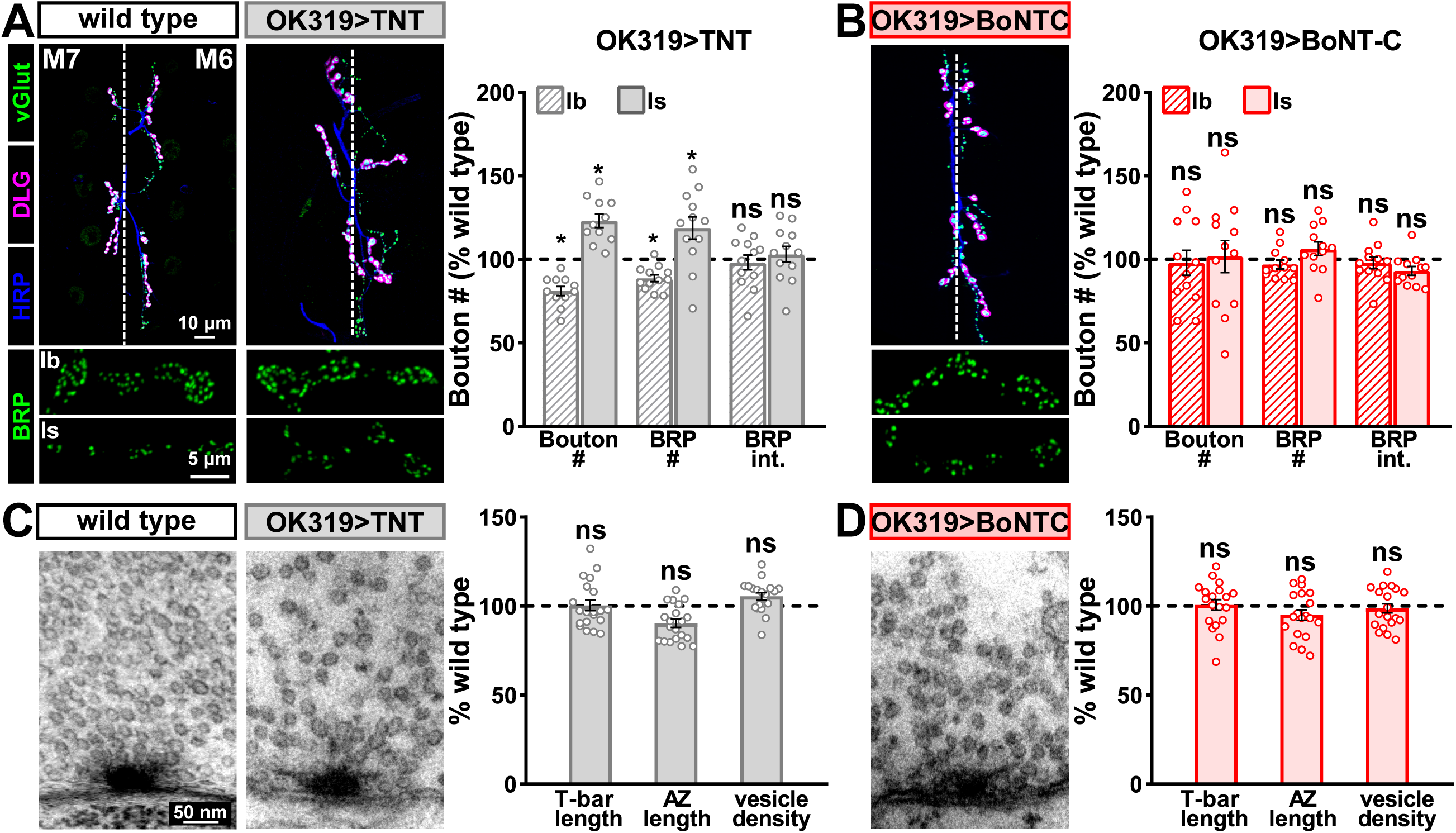
BoNT-C expression does not perturb presynaptic growth or structure. **(A)** Representative muscle 6 NMJ images immunostained with anti-vGlut, -DLG, and -HRP at wild type and NMJs expressing TNT; BRP immunostaining in individual boutons is shown below. Right: Quantification of bouton number, BRP number/NMJ, and mean BRP fluorescence intensity at TNT NMJs of MN-Ib or -Is expressing normalized to wild-type values. **(B)** Representative images and quantification of NMJs silenced by BoNT-C expression as described in (A). Note that in contrast to TNT, BoNT-C expression does not change bouton or BRP numbers at NMJs of either MN-Ib or -Is. **(C)** Representative electron micrographs of wild-type and TNT NMJs showing synaptic vesicles and active zone structures. Right: Quantification of T-bar length (µm), active zone length (µm) and synaptic vesicle density (#/µm^2^) normalized to wild-type values in the indicated genotypes. **(D)** Representative electron micrographs and analysis of BoNT-C NMJs as presented in (C). Note that no significant differences are observed compared to wild-type values. Error bars indicate ±SEM. *P<0.05; ns, not significant. Absolute values for normalized data and additional statical details are summarized in Table S2.

Finally, we determined whether BoNT-C expression in MNs altered postsynaptic structure at the NMJ. The postsynaptic glutamate receptors at the NMJ assemble as heterotetramers containing three essential subunits (GluRIII, GluRIID, and GluRIIE) and either a GluRIIA or GluRIIB subunit (Han et al., 2015; Qin et al., 2005). GluRIIA-containing receptors drive most of the synaptic currents due to the rapid desensitization of GluRIIB-containing subtypes (Diantonio et al., 1999; Han et al., 2015). Differences in GluR composition have been reported at postsynaptic compartments of MN-Is and MN-Ib, where GluRIIA-type receptors localize more centrally within GluR fields and are more abundant at MN-Ib postsynaptic compartments relative to MN-Is (Akbergenova et al., 2018; Marrus et al., 2004; Schmid et al., 2008). It has been speculated that tonic vs phasic patterns of activity at MN-Is and MN-Ib may orchestrate these differences in GluR composition at postsynaptic receptive fields (Aponte-Santiago and Littleton, 2020). We used synaptic silencing by BoNT-C as an opportunity to test this possibility. At wild-type postsynaptic compartments, we observed the lower GluRIIA intensity characteristic at MN-Is NMJs relative to MN-Ib (Fig. 4A,B). We also found the characteristic enrichment of GluRIIA at centers of receptive fields, with GluRIIB distributed in the periphery (Fig. 4C). Interestingly, these same input-specific patterns of GluR composition and spatial localization were observed following BoNT-C expression, where no glutamate is emitted from presynaptic terminals (Fig. 4D-F). These results demonstrate two important points. First, neurotransmitter release is not required for synaptic assembly and alignment at the *Drosophila* NMJ, nor for their maintenance during growth and elaboration, including the specialization of postsynaptic GluR fields. Second, patterns of tonic vs phasic transmission do not sculpt the final composition of postsynaptic receptive fields.

**Figure 4:**
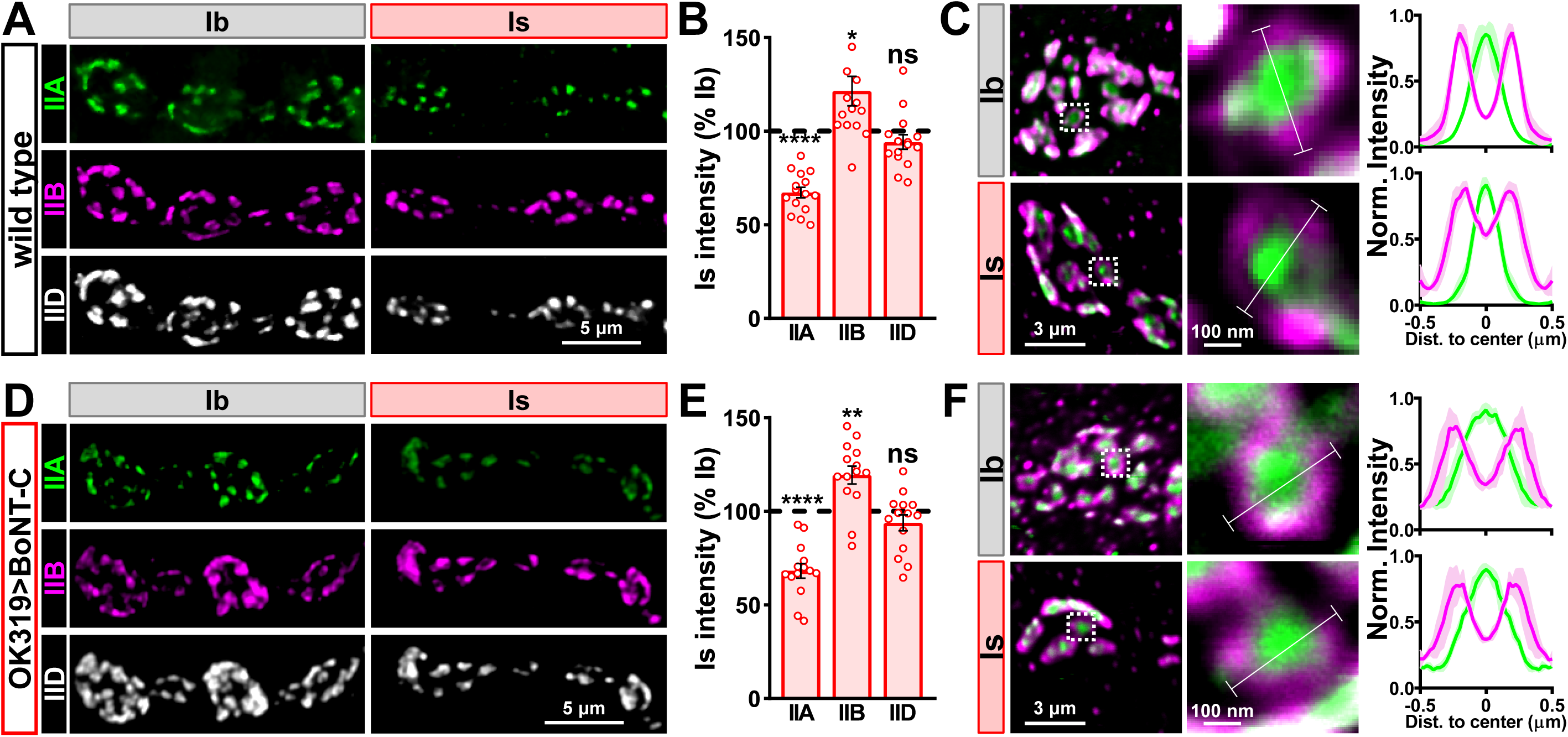
Tonic vs phasic activity patterns do not specialize postsynaptic GluR fields. **(A)** Representative images of boutons from MN-Ib and -Is NMJs immunostained with anti-GluRIIA, - GluRIIB and -GluRIID in wild type. **(B)** Quantification of the mean fluorescence intensity of each GluR subunit at MN-Is NMJs normalized as a percentage of those at MN-Ib. MN-Is NMJs exhibit a significant decrease in GluRIIA-containing GluRs, an increase in GluRIIB-containing GluRs, and no significant change in total GluR levels (indicated by the common subunit GluRIID) compared to GluR fields at MN-Ib NMJs. **(C)** High magnification image of individual NMJs imaged as in (A). Averaged fluorescence line profiles show GluRIIA and GluRIIB normalized to peak fluorescence values across 10 receptor fields in MN-Ib or -Is NMJs. Note that peak fluorescence of GluRIIA is at the center of the GluR field, while peak fluorescence of GluRIIB is located more peripherally. The white line indicates the line profile ROI. **(D-F)** Similar analysis of MN-Ib and -Is NMJs silenced by with BoNT-C expression (OK319>BoNT-C). Similar GluR levels and localizations are observed, indicating that tonic vs phasic patterns of transmission, and indeed glutamate release itself, is not required to establish the input-specific specialization of GluR fields. Error bars indicate ±SEM. ****P<0.0001; **P<0.01; *P<0.05; ns, not significant. Absolute values for normalized data and additional statistical details are summarized in Table S2.

### Input-specific silencing by BoNT-C does not induce heterosynaptic structural plasticity

Next, we sought to determine whether selective silencing of neurotransmitter release at MN-Is or MN-Ib by BoNT-C induced heterosynaptic structural plasticity. Previous studies have found heterosynaptic changes in synaptic growth following genetic ablation or TNT expression (Aponte-Santiago et al., 2020; Wang et al., 2021). First, we engineered the newest and fastest genetically encoded Ca^2+^ sensor (GCaMP8f) into a previous generation postsynaptic sensor (SynapGCaMP6f; (Newman et al., 2017)) to visualize input-specific elimination of synaptic transmission the larval NMJ. When BoNT-C is expressed in MN-Is only, Ca^2+^ imaging using SynapGCaMP8f revealed the complete elimination of both miniature and evoked transmission at MN-Is NMJs, while spontaneous and evoked transmission was unperturbed at MN-Ib NMJs (Fig. 5A-C). Conversely, selective BoNT-C expression in MN-Ib eliminated transmission from MN-Ib NMJs without impacting transmission from MN-Is (Fig. 5D-F). Thus, Ca^2+^ imaging using SynapGCaMP8f confirmed that selective BoNT-C expression can induce input-specific synaptic silencing.

**Figure 5:**
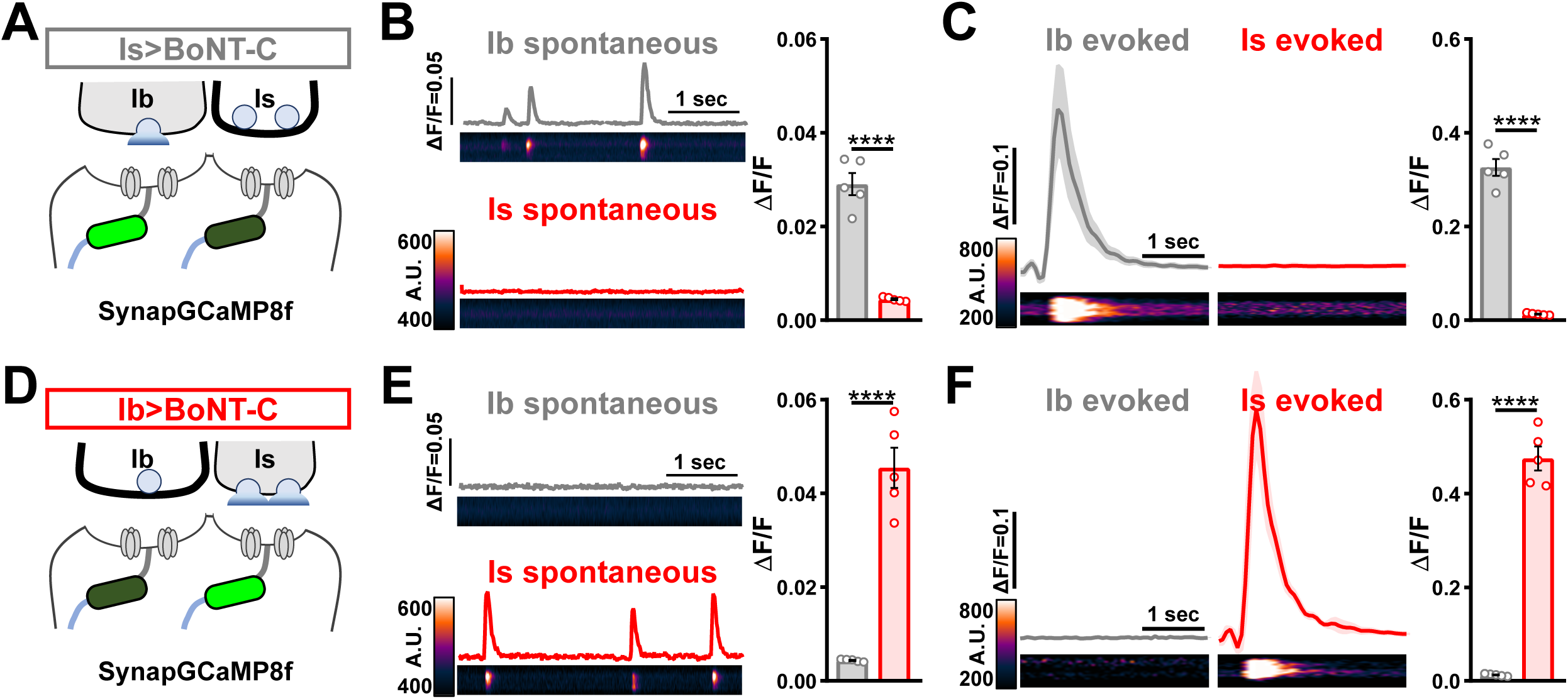
Postsynaptic Ca^2+^ signals are selectively eliminated by input-specific BoNT-C expression. **(A)** Schematic depicting silencing of MN-Is transmission by selective expression of BoNT-C in MN-Is. The postsynaptic Ca^2+^ indicator SynapGCaMP8f is indicated in the postsynaptic compartment, which should receive transmission only at MN-Ib NMJs. **(B)** Representative images and traces of spontaneous postsynaptic Ca^2+^ transients at MN-Ib and -Is NMJs using the new postsynaptic GCaMP8f indicator in Is>BoNT-C (*w*;*MHC>GCaMP8f*/*+*;*R29E07-GAL4*/*UAS-BoNT-C*). Right: Quantification of averaged transients, confirming that synaptic Ca^2+^ events are selectively eliminated at MN-Is NMJs but intact at MN-Ib. **(C)** Averaged traces and images of evoked postsynaptic Ca^2+^ transients at MN-Ib and -Is NMJs. Right: Quantification of averaged evoked transients, confirming that evoked synaptic Ca^2+^ events are selectively eliminated at MN-Is NMJs but intact at MN-Ib. **(D)** Schematic depicting silencing of MN-Ib transmission by selective expression of BoNT-C in MN-Ib. **(E-F)** Similar analysis as shown in (B-C) at MN-Ib silenced NMJs (Ib>BoNT-C; *w*;*MHC>GCaMP8f* /+;*dHB9-GAL4*/*UAS-BoNT-C*). Synaptic Ca^2+^ events are selectively eliminated at MN-Ib NMJs but intact at MN-Is. Error bars indicate ±SEM. ****P<0.0001. Additional statistical details are summarized in Table S2.

We then examined synaptic growth by counting Is vs Ib bouton numbers following input-specific expression of BoNT-C, TNT, and rpr.hid. When BoNT-C is expressed in MN-Is, no changes in synaptic growth in either MN-Is or MN-Ib was observed compared to wild type (Fig. 6A,B), demonstrating no heterosynaptic structural plasticity is induced by input-specific BoNT-C expression. Similarly, TNT expression in MN-Is increased Is bouton numbers, as expected (see Fig. 3), but no changes in Ib boutons numbers were observed (Fig. 6A,C), suggesting miniature-only transmission does not induce heterosynaptic structural changes. Finally, MN-Is ablation by rpr.hid expression fully ablated MN-Is inputs, while inducing a compensatory heterosynaptic increase in Ib bouton number (Fig. 6A,D), as previously observed at other larval NMJs (Aponte-Santiago et al., 2020; Wang et al., 2021). Parallel experiments driving BoNT-C, TNT, and rpr.hid transgenes in MN-Ib led to similar results as shown for MN-Is, with intrinsic but not heterosynaptic changes induced by *TNT* expression, ablation of both Is and Ib by rpr.hid expression, and no intrinsic or heterosynaptic structural plasticity induced by BoNT-C expression (Fig. 6E-H).

**Figure 6:**
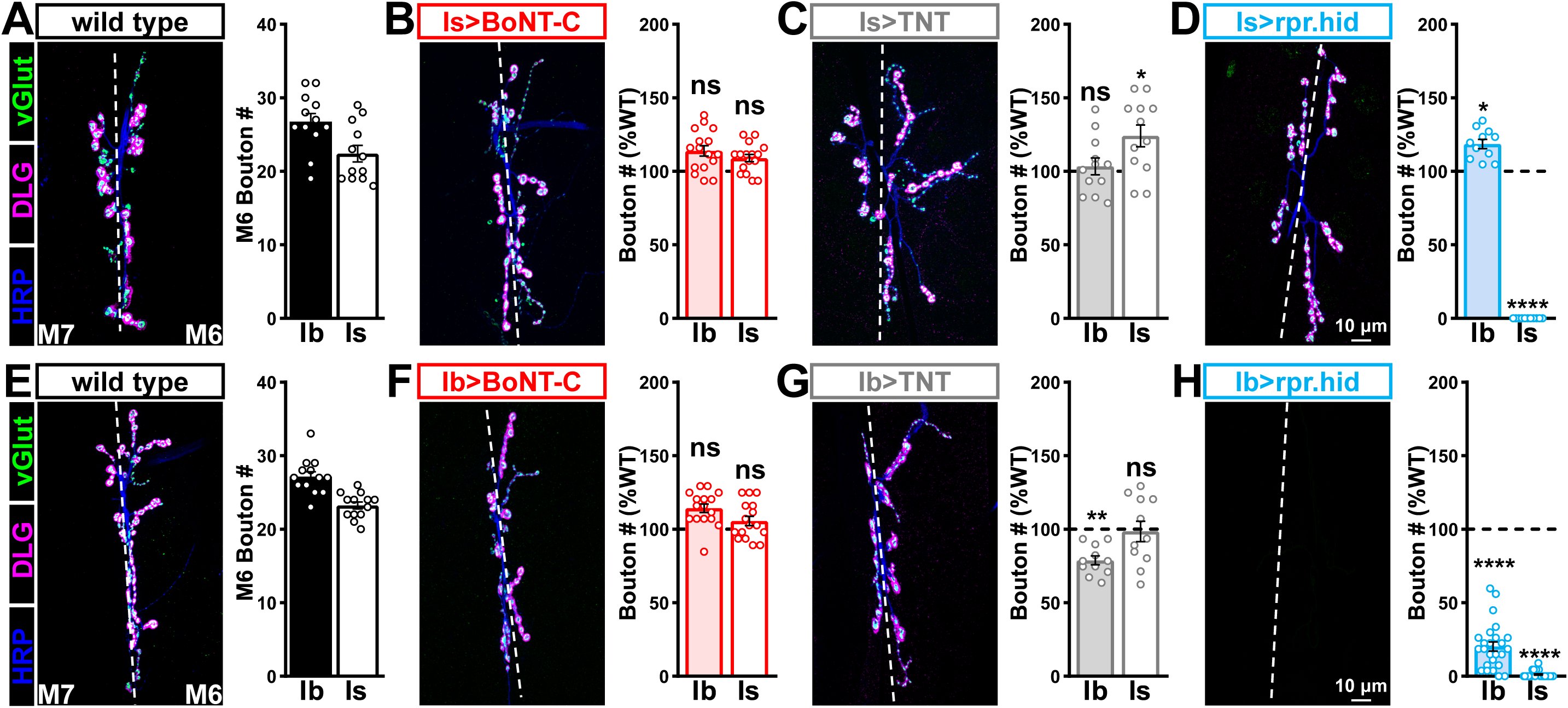
Selective silencing by BoNT-C does not induce heterosynaptic structural plasticity. **(A-D)** Representative NMJ images immunostained with anti-vGlut, -DLG, and -HRP in wild type (A); selective silencing of MN-Is (Is>BoNT-C; *w*;+;*R29E07-GAL4*/*UAS-BoNT-C*) (B); TNT expression in MN-Is (Is>TNT; *w*;*UAS-TNT*/+;*R29E07-GAL4*/+) (C); and ablation of MN-Is (Is>rpr.hid; *rpr.hid*/*+*;*R29E07-GAL4*/+) (D). Quantification of MN-Ib and MN-Is bouton number in wild type and in each condition normalized to wild-type values is shown to the right. Note that while no heterosynaptic structural plasticity is observed in bouton number Is>BoNT-C and Is>TNT, a compensatory heterosynaptic increase is found in Is>rpr.hid. **(E-H)** Similar NMJ images and quantification as shown in (A-D) but with MN-Ib expression of BoNT-C (Ib>BoNT-C; *w*;*+*;*dHB9-GAL4*/*UAS-BoNT-C*), -TNT (Ib>TNT; *w*;*UAS-TNT*/+;*dHB9-GAL4*/+), or -rpr.hid (Ib>rpr.hid; *rpr.hid*/+;*dHB9-GAL4*/+). Error bars indicate ±SEM. ****P<0.0001; ***P<0.001; *P<0.05; ns, not significant. Absolute values for normalized data and additional statistical details are summarized in Table S2.

It has been suggested that heterosynaptic plasticity may be distinctly expressed at different NMJs (Wang et al., 2021). We therefore examined synaptic growth at NMJs innervating muscles 12, 13, or 4 following TNT, rpr.hid, or BoNT-C expression in the Is MN innervating each of these muscles. However, intrinsic and heterosynaptic properties were similar at these other NMJs compared to our findings at muscle 6/7 (Table S2). Thus, while physical loss of MN-Is induced increased synaptic growth at MN-Ib, selective synaptic silencing by BoNT-C did not induce heterosynaptic structural plasticity. This reveals an important property about heterosynaptic structural plasticity at co-innervated muscles in *Drosophila*: No heterosynaptic structural plasticity is observed when the motor neuron is physically present but functionally silent.

### Selective silencing by BoNT-C fully reconstitutes wild type NMJ physiology

Complete neurotransmission from MN-Is and MN-Ib has never been electrophysiologically separated. Although input-specific silencing by BoNT-C does not induce heterosynaptic structural plasticity, it is possible that heterosynaptic functional adaptations might be imparted. Alternatively, input-specific silencing by BoNT-C may not induce functional changes in neurotransmission at the convergent input. We sought to distinguish between these possibilities. In standard electrophysiological recordings from wild-type NMJs, miniature transmission is an undefined blend of spontaneous events originating from both MN-Is and MN-Ib inputs, and evoked amplitudes reflect a nebulous composite of transmission from both motor inputs that does not reflect the behavior of any synapse that actually exists. We reasoned that if wild type physiology could be fully reconstituted after electrophysiologic isolation of both miniature and evoked transmission from MN-Is and MN-Ib, this would confirm that no heterosynaptic functional plasticity is induced by input-specific BoNT-C expression. However, if wild-type physiology was not fully reconstituted, this would indicate that heterosynaptic functional plasticity was provoked by input-specific BoNT-C silencing.

We first compared miniature transmission at wild type (Ib+Is), Ib only (Is>BoNT-C), and Is only (Ib>BoNT-C) NMJs. As expected, mEPSP frequency was reduced at both Ib and Is only compared to wild type (Fig. 7A,B). Importantly, substantial differences in mEPSP amplitude were observed at Is vs Ib, where miniature events were over 70% increased at MN-Is NMJs compared to MN-Ib (1.34 mV ±0.04 vs 0.77 mV ±0.02; Fig. 7A,C), as predicted by previous studies (Karunanithi et al., 2002; Newman et al., 2017). This highlights the inaccurate quantal sizes reported in previous studies where transmission from each input was blended, and averaged miniature events were used to estimate quantal content at composite (Ib+Is) evoked amplitudes or with selective optogenetic stimulation (Genç and Davis, 2019; Sauvola et al., 2021). To determine whether wild-type miniature physiology could be reconstituted using the input-specific BoNT-C data, we first summed the Ib only and Is only miniature frequencies together, which resulted in a 3.54 Hz frequency that was statistically indistinguishable to the wild-type value (Fig. 7B). Similarly, we combined the weighted average of mEPSP amplitudes from Ib only and Is only events, which provided a value of 1.00 mV ±0.01, statistically similar to the wild-type value (0.998 ±0.001; Fig. 7C). Thus, a reductionist analysis of miniature transmission from selectively silenced MN-Ib and MN-Is NMJs recapitulates the physiology observed at co-innervated muscles.

**Figure 7:**
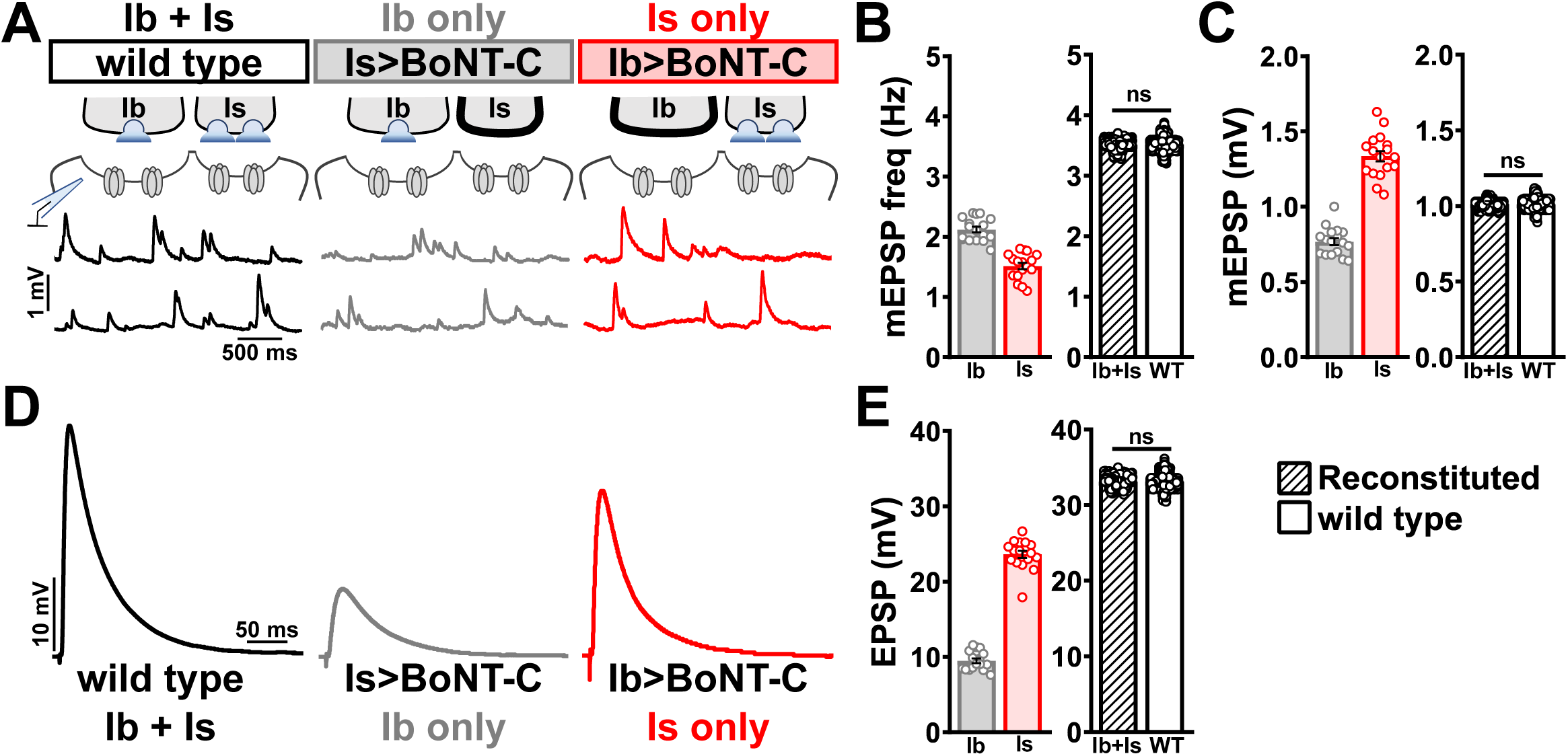
Selective silencing by BoNT-C fully reconstitutes wild type NMJ physiology. **(A)** Schematic and representative mEPSP traces of recordings from wild type (“Ib+Is”) or input-specific silencing of MN-Is (“Ib only”) or MN-Ib (“Is only”) by BoNT-C expression. **(B)** Quantification of mEPSP frequency in Ib only and Is only. A simple addition of these values (reconstituted) recapitulates the observed blended values observed in wild type. (C) Quantification of mEPSP amplitude in Ib only and Is only. Note the substantial input-specific difference in mEPSP amplitude. A weighted average of these values from Ib and Is (reconstituted) fully recapitulates the average mEPSP amplitude values observed from recordings of wild-type NMJs. Reconstituted data sets were acquired by bootstrapping and resampling from the seed data sets (see methods for additional details). **(D)** Representative EPSP traces from recordings wild type, Ib only, and Is only NMJs. **(E)** Quantification of EPSP amplitude from Ib only and Is only NMJs. A simple addition of these values (reconstituted) fully recapitulates the blended EPSP values obtained from wild-type recordings. Error bars indicate ±SEM. ns, not significant. Additional statistical details are summarized in Table S2.

Next, we examined evoked EPSP events from composite (Ib+Is) or isolated (Ib or Is only) NMJs. At 0.4 mM Ca^2+^ saline, evoked transmission from MN-Is is over two-fold higher than MN-Ib (23.65 ±0.48 vs 9.50 ±0.30; Fig. 7D,E). When we summed EPSP amplitude from paired MN-Ib and -Is inputs, we obtained EPSP values not statistically different from the composite values (33.10 ±0.03 vs 32.60 ±2.19; Fig. 7D,E), suggesting no heterosynaptic functional plasticity in evoked physiology is induced by selective BoNT-C expression. Finally, we were able to obtain an accurate quantal content value of synaptic vesicle release at MN-Ib and MN-Is, extracted from the disambiguated miniature and evoked transmission. Given the large quantal size of transmission from MN-Is, due to enhanced vesicle size (Karunanithi et al., 2002; Newman et al., 2017), less quantal content was observed than would be expected from the averaged quantal size, while the converse is true for transmission from MN-Ib (Table S2). Together, these data demonstrate that composite transmission from co-innervated muscles can be fully reconstituted from isolated MN-Ib and MN-Is transmission enabled by selective BoNT-C transmission. Further, this finding underscores that no heterosynaptic structural or functional plasticity is induced when MN-Is or MN-Ib innervation is physically present but functionally silent.

### Heterosynaptic functional plasticity is only induced by physical loss of the convergent input

In our final set of experiments, we sought to clarify whether heterosynaptic functional plasticity, like structural plasticity, could be induced through selective expression of TNT and/or rpr.hid. Previous studies have suggested possible changes in neurotransmission (Aponte-Santiago et al., 2020; Wang et al., 2021), but the inability to isolate input-specific baseline miniature and evoked transmission obfuscated to what extent heterosynaptic functional plasticity was induced. Using selective expression of BoNT-C to isolate baseline MN-Ib or MN-Is transmission, we then compared the physiology at either input following TNT or rpr.hid expression. When TNT is expressed in MN-Is, we observed similar EPSP amplitudes compared to Ib only values determined by BoNT-C expression in MN-Is (Fig. 8A), as well as the expected increase in miniature frequency and amplitude due to persistent miniature activity from Is>TNT (Fig. 8A). This indicates that synaptic silencing of evoked release from MN-Is using TNT does not induce heterosynaptic functional plasticity. However, while ablation of MN-Is by rpr.hid expression eliminated mEPSP events from Is, mirroring Is>BoNT-C values (Fig. 8B), an adaptive enhancement in heterosynaptic evoked transmission was observed (Fig. 8B). Parallel experiments selectively expressing TNT in MN-Ib revealed no change in evoked transmission (Fig. 8C), consistent with TNT expression in MN-Is. Similarly, selective ablation of MN-Ib by Ib>rpr.hid expression eliminated most transmission (Fig. 8D) due to loss of both MN-Ib and -Is inputs (Fig. S4).

**Figure 8:**
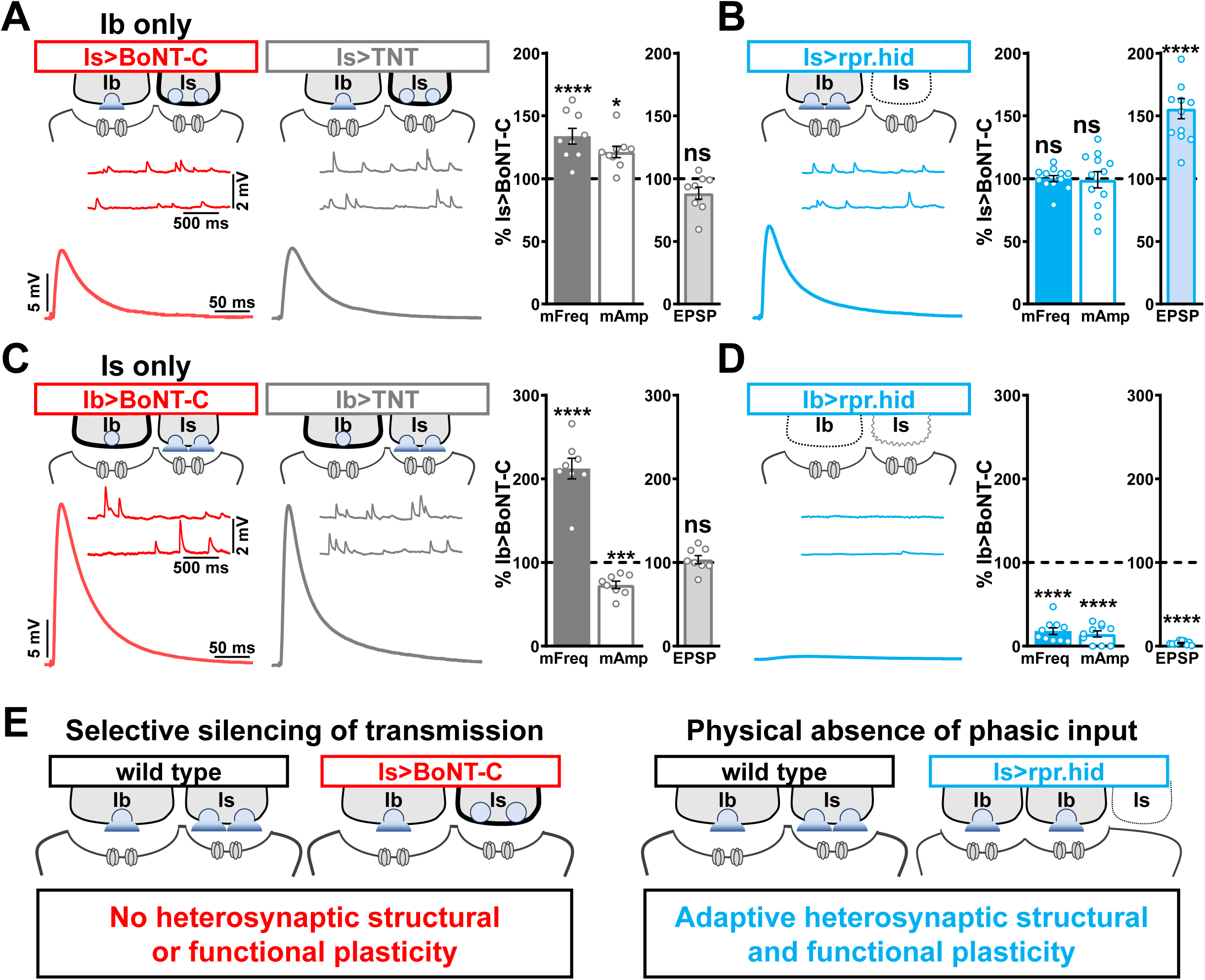
Selective silencing by BoNT-C does not induce heterosynaptic functional plasticity. **(A,B)** Schematic and representative electrophysiological traces of Ib only NMJs (Is>BoNT-C) compared to Is>TNT (A) and Is>rpr.hid (B). Right: Quantification of mEPSP frequency and amplitude as well as EPSP amplitude normalized to Is>BoNT-C values in the indicated genotypes. Note that in Is>TNT, mEPSP values are significantly increased, as expected, due to persistent mEPSP release from MN-Is, while no significant change is observed in EPSP amplitude. In contrast, no difference in mEPSP values are observed following MN-Is ablation by *rpr.hid* expression, while an apparent adaptive heterosynaptic increase in EPSP amplitude is observed from the remaining MN-Ib. **(C,D)** Schematic and representative traces of Is only NMJs (Ib>BoNT-C) compared to Ib>TNT (C) and Ib>rpr.hid (D). Right: Quantification of mEPSP frequency and amplitude as well as EPSP amplitude normalized to Ib>BoNT-C values in the indicated genotypes. Note that in this case, the expected changes in mEPSP values but no significance difference in EPSP values are observed in Ib>TNT. However, mEPSP and EPSP values are essentially eliminated in Ib>rpr.hid due to ablation of both MN-Ib and -Is inputs. **(E,F)** Schematics summarizing heterosynaptic plasticity at the *Drosophila* NMJ. When transmission is silenced by BoNT-C but both motor inputs are physically present, no heterosynaptic structural or functional plasticity is elicited (E). However, when a motor input is physically absent, adaptive heterosynaptic structural and functional plasticity is induced (F). Error bars indicate ±SEM. ****P<0.0001; ***P<0.001; ns, not significant. Absolute values for normalized data and additional statistical details are summarized in Table S2.

Together, these results illustrate important heterosynaptic plasticity rules between tonic and phasic neurons (Fig. 8E,F). When neurotransmission is silenced from one input, no heterosynaptic structural or functional plasticity is induced. However, physical loss of one input provokes an adaptive increase in synaptic strength due to enhanced synapse number and neurotransmitter release from the convergent input.

## DISCUSSION

By screening a variety of botulinum neurotoxins, we identified BoNT-C to silence all neurotransmission when expressed in *Drosophila* MNs. Crucially, BoNT-C expression does not impair NMJ growth or synaptic structure and can isolate neurotransmission from phasic MN-Is and tonic MN-Ib when selectively expressed in either MN. Finally, no heterosynaptic structural or functional plasticity is induced at the convergent input following selective BoNT-C expression, revealing that heterosynaptic adaptive plasticity requires physical loss of the motor input. Thus, BoNT-C provides a powerful new approach to accurately disambiguate tonic vs phasic neurotransmission at the *Drosophila* NMJ and provides a foundation from which to understand the induction mechanisms of heterosynaptic plasticity at this model glutamatergic synapse.

### BoNT-C as a tool for effectively silencing neurotransmission

Neurotoxins that target SNARE complexes have been utilized in neuroscience research for decades. In *Drosophila*, transgenic control of tetanus toxin light chain (TNT) expression was established decades ago (Sweeney et al., 1995) and widely used since to inhibit transmission in both peripheral and central neurons. However, while TNT expression eliminates evoked neurotransmission, miniature transmission persists (Baines et al., 2001; Choi et al., 2014; Goel and Dickman, 2018), as also observed in *n-syb* mutations (Deitcher et al., 1998), the target of TNT. This persistence of miniature transmission following TNT intoxication of MNs was exploited to explore the functions of spontaneous neurotransmission in synaptic growth and development at the *Drosophila* NMJ (Choi et al., 2014) and the proportion of postsynaptic Ca^2+^ influx driven by evoked released at tonic vs phasic NMJs (Newman et al., 2017). However, despite the identification of GAL4 drivers specific for tonic vs phasic MNs, two major limitations rendered TNT a sub-optimal approach to isolate input-specific transmission at the fly NMJ. First, the persistence of miniature transmission obfuscates the determination of accurate quantal content values from either MN, particularly given the large difference in quantal size between the two inputs (Fig. 7). In addition, TNT expression alters synaptic growth at MN terminals (Fig. 3 and (Li et al., 2018)), and heterosynaptic plasticity was reported with input-specific TNT expression (Aponte-Santiago et al., 2020). Interestingly, while we did not observe heterosynaptic structural or functional plasticity following input-specific TNT expression at the muscle 6/7 NMJ, accurate evoked EPSP amplitudes were observed (Fig. 8), suggesting that input-specific TNT expression can alter synaptic growth and block evoked transmission without inducing any apparent changes at the convergent input.

The isolation of BoNT-C was fortunate given the variable activity and toxicity previously reported with BoNTs. In the *Drosophila* visual system, BoNT transgene expression led Backhaus to variable inhibition of transmission and substrate cleavage where, for example, BoNT-A only partially cleaved SNAP-25 and Syntaxin (Backhaus et al., 2016). Similarly, we also observed variable activity between independent transgenic inserts of BoNT-E, where expression of some transgenes appeared to make animals less healthy than others (see methods). In addition, expression of TNT or BoNT can lead to neurodegeneration (Berliocchi et al., 2005; Peng et al., 2013; Zhao et al., 2010). Although we cannot rule out that BoNT-C may induce toxicity when expressed in central neurons or for longer durations such as in adult stages, we find no evidence for toxicity or altered health when BoNT-C is expressed in MNs throughout larval development. This is likely due to BoNT-C exhibiting specific cleavage activity for *Drosophila* Syntaxin (Backhaus et al., 2016) as well as a fortuitous genomic insertion that ensures moderate expression of the transgene.

Several important properties of synaptic structure and function in tonic vs phasic MNs have been confirmed and extended by selective BoNT-C silencing. First, phasic NMJs are responsible for about twice the postsynaptic depolarization compared to tonic, at least with regard to single stimuli under the conditions used in our study. However, despite this obvious difference in synaptic strength, quantal content values released between MN-Is vs -Ib are more similar than would be apparent based on the substantial difference in quantal size. Second, tonic vs phasic firing patterns do not determine the specialized abundance and localization of postsynaptic glutamate receptive fields. Third, synaptic morphogenesis and structure does not require vesicular neurotransmitter release at glutamatergic synapses, as BoNT-C NMJs appear indistinguishable from wild type. Although several studies have shown neurotransmitter release is not necessary for the initial establishment of synapses (Banerjee et al., 2022; Sando et al., 2017; Sigler et al., 2017; Varoqueaux et al., 2002), whether miniature transmission was necessary for clustering of glutamate receptors at the fly NMJ has provoked some controversy (Otsu and Murphy, 2003; Saitoe et al., 2002a, 2002b; Verstreken and Bellen, 2002). It will be of significant interest in future studies to now determine the full electrophysiological properties of tonic vs phasic neurons, including vesicle pools, short term plasticity, and release probability across various extracellular Ca^2+^ conditions. Furthermore, to what extent synaptic components exhibit specialized functions at tonic vs phasic MNs, as was recently shown for *tomosyn* (Sauvola et al., 2021), will also be an exciting area to leverage BoNT-C silencing going forward. Input-specific silencing by BoNT-C now provides a foundation to elucidate fundamental electrophysiological differences between tonic and phasic neuronal subtypes and, importantly, to re-evaluate specialized functions of *Drosophila* synaptic genes in which initial characterizations relied on blended transmission from both inputs.

### Insights into synapse development and heterosynaptic plasticity revealed by BoNT-C

Physical loss of tonic or phasic motor inputs appears to be the key inductive process capable of inducing adaptive heterosynaptic plasticity. Input-specific silencing of neurotransmission by BoNT-C, or even blockade of evoked release only by TNT, failed to elicit any structural or functional changes in the convergent input at the NMJs we assayed (muscle 6/7, 12/13, and 4). Genetic ablation of MN-Ib largely eliminated tonic inputs, as expected, and completely prevented phasic innervation at the convergent muscle 6/7 NMJ. This dramatic change is in contrast to more subtle differences in MN-Is innervation following MN-Ib ablation at the muscle 1 NMJ (Aponte-Santiago et al., 2020). Indeed, we found no evidence that phasic innervation ever occurred when tonic MNs were ablated, with no remnants of phasic postsynaptic structures remaining at the muscle target, while other muscle targets innervated by the same phasic input appeared largely unchanged. This is in contrast to models of neuromuscular degeneration, in which “synaptic footprints” of postsynaptic structures remain after initial synaptogenesis and growth followed by MN retraction (Eaton et al., 2002; Perry et al., 2017). The failure of phasic innervation following tonic ablation suggests that during development, the tonic MN provides necessary “instructive” cues for proper phasic innervation, akin to pioneer axons important for axon guidance (Araújo and Tear, 2003; Raper and Mason, 2010). Tonic MN-Ib serving as guidance cues for phasic innervation may make sense given that each muscle target receives tonic innervation from a single MN, while phasic MNs innervate groups of multiple muscles (Hoang and Chiba, 2001).

Conversely, ablation of phasic MN-Is inputs was the only condition in which we found evidence for adaptive structural and functional heterosynaptic plasticity at tonic MN-Ib inputs. It was in this condition where clear evidence for adaptive heterosynaptic structural plasticity was also reported in previous studies (Wang et al., 2021). Importantly, BoNT-C silencing establishes that loss of neurotransmission from the phasic input *per se* is not sufficient to induce any adaptive heterosynaptic plasticity. Rather, physical loss of the phasic input must provide an instructive cue, perhaps through signaling from peripheral glia and/or the common muscle target, that elicits adaptive plasticity at the remaining tonic input.

It would be appealing if loss of transmission from one input could induce a homeostatic adjustment at the convergent input that maintains stable postsynaptic excitation. Clearly this is not the case at the *Drosophila* NMJ. BoNT-C expression in the tonic input depresses transmission by ∼1/3, while loss of phasic transmission reduces NMJ strength by ∼2/3, with no evidence of heterosynaptic plasticity observed. In our study and in other cases where heterosynaptic plasticity has been reported (Aponte-Santiago et al., 2020; Wang et al., 2021), the seemingly adaptive changes in synaptic growth or function are quite subtle, nowhere near the levels that would be necessary to restore basal levels of transmission. For example, total bouton number remained highly reduced following phasic MN-Is ablation (wild type: 52±1.5; Is>rpr.hid: 32±0.8 bouton numbers), with a corresponding reduction in synaptic strength (wild type: 32.6±2.2 mV; Is>rpr.hid: 14.2±0.7 mV EPSP values). However, while loss of transmission or innervation to a particular target muscle does not induce homeostatic plasticity, a converse manipulation, in which innervation is biased between adjacent muscle targets, does elicit two distinct forms of target-specific homeostatic plasticity that stabilizes synaptic strength (Davis and Goodman, 1998; Goel et al., 2020). Hyper-innervation of both tonic and phasic inputs on muscle 6 homeostatically reduces release probability from both inputs without any obvious postsynaptic changes (Davis and Goodman, 1998; Goel et al., 2020). In contrast, hypo-innervation of both tonic and phasic inputs on the adjacent muscle 7 leads to no functional changes in the neurons, while a homeostatic enhancement in postsynaptic glutamate receptor abundance compensates for reduced transmitter release to restore synaptic strength (Goel et al., 2020). Illuminating the complex synaptic dialogue between pre- and post-synaptic compartments within and between common targets will clarify how adaptive and homeostatic plasticity mechanisms are engaged in at NMJs and in motor circuits in general.

## MATERIALS AND METHODS

### Fly stocks

Experimental flies were raised at 25°C on standard molasses food. The *w^1118^* strain was used as the wild type control unless otherwise noted, as this is the genetic background for which all genotypes are bred. For optogenetic experiments, flies were raised in consistent dark conditions on standard food supplemented with 500 µM all-trans-retinal (#R2500, Sigma-Aldrich). Second-instar larvae were then transferred to fresh food containing 500 µM all-trans-retinal. The following fly stocks were used: *OK6-GAL4* (Aberle et al., 2002), *UAS-rpr.hid* (Zhou et al., 1997), *OK319-GAL4* (Sweeney et al., 1995), *vGlut^SS1^* (Sherer et al., 2020), *B3RT-vGlut-B3RT* (Sherer et al., 2020), *UAS-B3* (Sherer et al., 2020). The following fly strains were generated in this study: *UAS-BoNT-A*, *UAS-BoNT-B*, *UAS-BoNT-C*, *UAS-BoNT-E*, SynapGCaMP8f (*MHC-CD8-GCaMP8f-sh*). All other stocks were obtained from Bloomington Drosophila Stock Center (BDSC): *UAS-ChR2^T159C^* (#58373), *dHb9-GAL4* (Ib-GAL4, #83004), *GMR27E09-GAL4* (Is-GAL4, #49227), *UAS-TNT* (#28838), *w^1118^* (#5905), *D42-GAL4* (#8816), *UAS-CD4::tdGFP* (#35839). Details of these and additional stocks and their sources are listed in Table S3.

### Molecular Biology

To generate UAS-BoNT-A, -B, - C, and -E transgenes, we cloned sequences encoding each BoNT light chain (from plasmids shared by Matt Kennedy, University of Colorado, USA) into the gateway vector donor plasmid (ThermoFisher, # K240020). We then transferred these sequences into the final *pUASt* destination vector from Carnegie Drosophila Gateway Vector Collection using the Gateway reaction kit (Thermofisher, #11791020). To generate SynapGCaMP8f (*MHC-CD8-GCaMP8f-Sh*), we obtained the SynapGCaMP6f transgenic construct (Newman et al., 2017) and replaced the sequence encoding GCaMP6f with a sequence encoding GCaMP8f (#162379, Addgene) using Gibson assembly. Transgenic stocks were inserted into *w^1118^* by Bestgene, Inc (Chino Hills, CA, USA) using P-element-mediated random insertion and subsequently mapped and balanced. Pilot crosses of various BoNT transgenes to various neural GAL4 drivers were first used to select the inserts that appeared to induce the most consistent lethality and used for further analysis.

### Electrophysiology

Electrophysiological recordings were performed as described (Kiragasi et al., 2020; Li et al., 2021) using modified hemolymph-like saline (HL-3) containing: 70mM NaCl, 5mM KCl, 10mM MgCl_2_, 10mM NaHCO_3_, 115mM Sucrose, 5mM Trehelose, 5mM HEPES, and 0.5mM CaCl_2_, pH 7.2, from cells with resting potentials between -60 and -75 mV and input resistances >6 MΩ. Recordings were performed on an Olympus BX61 WI microscope using a 40x/0.80 NA water-dipping objective and acquired using an Axoclamp 900A amplifier, Digidata 1440A acquisition system and pClamp 10.5 software (Molecular Devices). Miniature excitatory postsynaptic potentials (mEPSPs) were recorded in the absence of any stimulation. Excitatory postsynaptic potentials (EPSPs) were recorded by delivering 20 electrical stimulations at 0.5 Hz with 0.5 msec duration to motor neurons using an ISO-Flex stimulus isolator (A.M.P.I.) with stimulus intensities set to avoid multiple EPSPs. All recordings were made on abdominal muscle 6 in segments A2 or A3 of third-instar larvae of both sexes. Data were analyzed using Clampfit (Molecular Devices), MiniΑnalysis (Synaptosoft), or Excel (Microsoft). Averaged mEPSP amplitude, mEPSP frequency, EPSP amplitude, and quantal content values were calculated for each genotype. Simulated data in Figure 7 of WT, Is and Ib were acquired by bootstrapping and resampling. EPSP, mEPSP and mEPSP frequency data were resampled 1000 times from the raw seed data set shown in the left panels in Figure 7B, C and E and Table S1. 1000 mean values were calculated from all the resampled data sets, followed by calculating EPSP amplitude, mEPSP frequency and mEPSP amplitude for paired Is + Ib using the following equation: *EPSP_(Is+Ib)_ = EPSP_(Is)_ + EPSP_(Ib)_*; *freq._(Is+Ib)_ = freq._(Is)_ + freq._(Is)_*; *mEPSP_(Is+Ib)_ = (mEPSP_(Is)_ x freq._(Is)_ + mEPSP_(Ib)_ x freq._(Is)_)/freq._(Is+Ib)_*.

### Ca^2+^ imaging and analysis

Third-instar larvae were dissected in ice-cold saline. Imaging was performed in modified HL-3 saline with 1.5 mM Ca^2+^ added using a Zeiss Examiner A1 widefield microscope equipped with a 63x/1.0 NA water immersion objective. NMJs on muscle 6 were imaged at a frequency of 100 fps (512 x 256 pixels) with a 470 nm LED light source (Thorlabs) using a PCO sCMOS4.2 camera. Spontaneous Ca^2+^ events were imaged at NMJs during 120 sec imaging sessions from at least two different larvae. Evoked Ca^2+^ events were induced by delivering 10 electrical stimulations at 0.5 Hz. Horizontal drifting was corrected using ImageJ plugins (Kang Li, 2008) and imaging data with severe muscle movements were rejected as described (Ding et al., 2019). Three ROIs were manually selected using the outer edge of terminal Ib boutons observed by baseline GCaMP signals with ImageJ (Rueden et al., 2017; Schindelin et al., 2012). Ib and Is boutons were defined by baseline GCaMP8f fluorescence levels, which are 2-3 fold higher at Ib NMJs compared to their Is counterparts at a particular muscle. Fluorescence intensities were measured as the mean intensity of all pixels in each individual ROI. ΔF for a spontaneous event was calculated by subtracting the baseline GCaMP fluorescence level F from the peak intensity of the GCaMP signal during each spontaneous event at a particular bouton as previously detailed (Li et al., 2021). Baseline GCaMP fluorescence was defined as the average fluorescence of 2 secs in each ROI without spontaneous events. ΔF/F was calculated by normalizing ΔF to baseline signal F. For each ROI under consideration, the spontaneous event ΔF/F value was averaged for all events in the 60 sec time range to obtain the mean quantal size for each bouton. Data analysis was performed with customized Jupyter Note codes.

### Optogenetics

Electrophysiology recording with optogenetics stimulation were performed with the same set up detailed above with a 40x/1.0 NA water immersion objective. Dissection and electrophysiological recordings were performed in the same modified HL-3 saline described above. To stimulate EPSP events, 0.5 msec light pulses of 470 nm were delivered from an LED driver (Thorlabs) at 0.5 Hz, triggered by a TTL pulse driven from a Digidata 1440 DAC programmed using pClamp 10.5 software (Molecular Devices). Light intensity was controlled by the LED driver to avoid multiple EPSP events. The power range were calibrated between 5 to 20 mW/cm^2^ under the objective.

### Immunocytochemistry

Third-instar larvae were dissected in ice cold 0 Ca^2+^ HL-3 and immunostained as described (Goel and Dickman, 2018). In brief, larvae were either fixed in Bouin’s fixative for 5 min (Sigma, HT10132-1L), 100% ice-cold ethanol for 5 min, or 4% paraformaldehyde (PFA) for 10 min. Larvae were then washed with PBS containing 0.1% Triton X-100 (PBST) for 30 min, blocked with 5% Normal Donkey Serum followed by overnight incubation in primary antibodies at 4°C. Preparations were then washed 3x in PBST, incubated in secondary antibodies for 2 hours, washed 3x in PBST, and equilibrated in 70% glycerol. Prior to imaging, samples were mounted in VectaShield (Vector Laboratories). Details of all antibodies, their source, dilution used, and references are listed in Table S3.

### Confocal imaging and analysis

Samples were imaged as described (Kikuma et al., 2019) using a Nikon A1R Resonant Scanning Confocal microscope equipped with NIS Elements software and a 100x APO 1.4NA oil immersion objective using separate channels with four laser lines (405 nm, 488 nm, 561 nm, and 647 nm). For fluorescence intensity quantifications of BRP, vGlut, GluRIIA, GluRIIB and GluRIID, z-stacks were obtained on the same day using identical gain and laser power settings with z-axis spacing between 0.15-0.20 µm for all genotypes within an individual experiment. Maximum intensity projections were utilized for quantitative image analysis using the general analysis toolkit of NIS Elements software. The fluorescence intensity levels of BRP, vGlut, GluRIIA, GluRIIB and GluRIID immunostaining were quantified by applying intensity thresholds and filters to binary layers in the 405 nm, 488 nm, or 561 nm channels. The mean intensity for each channel was quantified by obtaining the average total fluorescence signal for each individual punctum and dividing this value by the puncta area. A mask was created around the HRP channel, used to define the neuronal membrane, and only puncta within this mask were analyzed to eliminate background signals. Boutons were defined as vGlut puncta, and DLG co-staining was used to define boutons at MN-Is vs -Ib NMJs. All measurements based on confocal images were taken from synapses acquired from at least six different animals.

### Electron microscopy

EM analysis was performed as described previously (Goel et al., 2019). Wandering third-instar larvae were dissected in Ca^2+^-free HL-3 and then fixed in 2.5% glutaraldehyde/0.1 M cacodylate buffer at 4°C. Larvae were then washed three times for 20 min in 0.1 M cacodylate buffer. The larval pelts were then placed in 1% osmium tetroxide/0.1M cacodylate buffer for 1 h at room temperature. After washing the larva twice with cacodylate and twice with water, larvae were then dehydrated in ethanol. Samples were cleared in propylene oxide and infiltrated with 50% Eponate 12 in propylene oxide overnight. The following day, samples were embedded in fresh Eponate 12. EM sections were obtained on a Morgagni 268 transmission electron microscope (FEI). NMJs were serial sectioned at a 60- to 70-nm thickness. The sections were mounted on Formvar-coated single slot grids and viewed at a 23,000 magnification and were recorded with a Megaview II CCD camera. Images were analyzed using the general analysis toolkit in the NIS Elements software and ImageJ software. Active zone area was measured by defining a circle area with a diameter of the length of the active zone dense at the center of T-bar structure. Synaptic vesicle density was analyzed by vesicle numbers normalized to active zone area.

### Statistical analysis

Data were analyzed using GraphPad Prism (version 7.0) or Microsoft Excel software (version 16.22). Sample values were tested for normality using the D’Agostino & Pearson omnibus normality test which determined that the assumption of normality of the sample distribution was not violated. Data were then compared using either a one-way ANOVA and tested for significance using a Tukey’s multiple comparison test or using an unpaired 2-tailed Student’s t-test with Welch’s correction. In all figures, error bars indicate ±SEM, with the following statistical significance: p<0.05 (*), p<0.01 (**), p<0.001 (***), p<0.0001 (****); ns=not significant. Additional statistics and sample number n values for all experiments are summarized in Tables S1 and S2.

## ACKNOWLEDGEMENTS

We thank Matt Kennedy (University of Colorado, Aurora, CO, USA) for sharing botulinum neurotoxin plasmids, Zachary Newman and Udi Issacof (UC Berkeley, Berkeley, CA, USA) for sharing the SynapGCaMP6f plasmid, Steve Stowers (Montana State University, Bozeman, MT, USA) for sharing the vGlut conditional knock out lines, and Greg Macleod (Florida Atlantic University, Jupiter, FL, USA) for important discussions on MN-Is and MN-Ib physiology and for comments on an earlier version of this manuscript. We acknowledge the Developmental Studies Hybridoma Bank (Iowa City, Iowa, USA) for antibodies used in this study and the Bloomington Drosophila Stock Center for fly stocks (NIH P40OD018537). We thank Landon Porter and Brian Leung for assistance in validating reagents. This work was supported by grants from the National Institutes of Health (NS091546 and NS111414) to DD.

## AUTHOR CONTRIBUTIONS

Electron microscopy was performed by CP and CB and analyzed by KH. Early BoNT studies were performed by PG. YH obtained and analyzed all other experimental data, with help from CC. The manuscript was written by YH and DD with feedback from the other authors.

## FIGURE LEGENDS

**Figure S1: Motor neuron-specific GAL4 expression at the *Drosophila* NMJ. (A)** Schematic of the Ib and Is MNs that innervate the indicated muscles. The pan-MN driver *OK6-GAL4* is expressed in all larval MNs. Right: Images of larval body wall muscles (labeled by Phalloidin) innervated by Ib and Is motor neurons (labeled with GFP; OK6>tdGFP: *w*;*OK6-GAL4*/*UAS-CD4::tdGFP*;*+*) and the post-synaptic density marker DLG to distinguish Ib vs Is NMJs. **(B)** Schematic and images of the Ib and Is MNs targeted by *OK319-GAL4*, which is expressed in two Is MNs and the two Ib MNs that innervate muscles 6/7 and 4 (OK319>tdGFP: *w*;*OK319-GAL4*/*UAS-CD4::tdGFP*;*+*). **(C)** Schematic and images of the two Ib MNs targeted by the MN-Ib-specific driver *dHB9-GAL4*, which is expressed in the three MN-Ibs that innervate muscles 6/7, 12, and 13 (Ib>tdGFP: *w*;*UAS-CD4::tdGFP*/+;*dHb9-GAL4*/*+*). **(D)** Schematic and images of the two Is motor neurons targeted by the MN-Is-specific driver *R27E09-GAL4*, which is expressed in both Is motor neurons (Is>tdGFP: *w*;*UAS-CD4::tdGFP*/+;*R29E07-GAL4*/*+*).

**Figure S2: vGlut expression persists despite conditional knock out in motor neurons. (A)** Schematic and representative mEPSP and EPSP traces illustrating transmission persists in conditional knock out of vGlut in MN-Ib or -Is. Genotypes: wild type (*w^1118^*); Is>vGlut^-/-^ (*w*;*vGlut^SS1^*/*B3RT-vGlut-B3RT*;*UAS-B3*/*R7E09-GAL4*); Ib>vGlut^-/-^ (*w*;*vGlut^SS1^*/*B3RT-vGlut-B3RT*;*UAS-B3*/*dHB9-GAL4*). **(B)** Quantification of mEPSP frequency, amplitude, EPSP, and quantal content in the indicated genotypes. **(C)** Representative images of MN-Ib and -Is NMJs immunostained with anti-vGlut, and the neuronal membrane marker HRP in the indicated genotypes. D42>vGlut^-/-^ (pan-motor neuron; *w*;*vGlut^SS1^*/*B3RT-vGlut-B3RT*;*UAS-B3*/*D42-GAL4*)>vGlut^-/-^. **(D)** Quantification of mean fluorescence intensity of vGlut in the indicated genotypes normalized to wild-type values. ****P<0.0001; *P<0.05; ns, not significant. Absolute values for normalized data and additional statistical details are summarized in Table S2.

**Figure S3: Electrophysiological differences between optogenetic and electrical stimulation at the *Drosophila* NMJ. (A)** Representative EPSP traces demonstrating that ChR expression in motor neurons reduces evoked transmission using either electrical or optical stimulation. Genotype: OK6>ChR2^T159C^ (*w*;*OK6-GAL4*/*UAS-ChR2^T159C^*;+). **(B)** Quantification of EPSP amplitude in the indicated genotypes and stimulation. **(C-F)** Representative EPSP traces and quantification showing that selective ChR expression in MN-Is (C,D) or -Ib (E,F) similarly reduces transmission compared to input-specific silencing by BoNT-C expression. Genotypes: Is>BoNT-C (*w*;+;*R29E07-GAL4*/*UAS-BoNT-C*); Ib>ChR2^T159C^ (*w*;*dHB9-GAL4*/*UAS-ChR2^T159C^*;*+*); Ib>BoNT-C (*w*;+;*dHB9-GAL4*/*UAS-BoNT-C*); Is>ChR2^T159C^ (*w*;*R29E07-GAL4*/*UAS-ChR2^T159C^*;*+*). ****P<0.0001; **P<0.01; *P<0.05. Additional statistical details are summarized in Table S2.

**Figure S4: Genetic ablation of MN-Ib abolishes innervation by MN-Is on the same target. (A)** Representative muscle 6/7 NMJ images showing MN-Is labeled by CD4::tdGFP (Is>CD4::tdGFP: *w*;*UAS-CD4::tdGFP*/+;*R29E07-GAL4*/*+*), co-stained with anti-DLG and - Phalloidin. **(B)** Representative NMJ images of MN-Is ablation by *rpr.hid* expression (Is>rpr.hid; *rpr.hid*/*+*;*+*;*R29E07-GAL4*/+) co-stained with anti-vGlut, -DLG and -HRP. Right: Quantification of MN-Is bouton number normalized to wild-type values in the indicated genotype, confirming full MN-Is ablation. **(C)** Representative NMJ images of MN-Ib labeled with GFP expression (Ib>CD4::tdGFP: *w*;*UAS-CD4::tdGFP*/+;*dHb9-GAL4*/*+*), co-stained with anti-DLG and - Phalloidin. **(D)** Representative NMJ images of wild type and MN-Ib ablation (Ib>rpr.hid; *rpr.hid*/*+*;*+*;*dHB9-GAL4*/+) immunostained with anti-vGlut, -DLG and -HRP, demonstrating that ablation of MN-Ib leads to loss of MN-Is innervation. Right: Quantification of MN-Ib bouton number normalized to wild-type values in the indicated genotypes. **(E-F)** Representative NMJ images of MN-Ib ablation illustrating two outcomes across 27 samples. Although MN-Is innervation was completely lost in all samples, 37% of NMJ samples exhibited a total loss of MN-Ib innervation as well (consistent with a failure to evoke an EPSP response before staining) (E). In contrast, 62% of cases were found to retain partial MN-Ib innervation (F). Quantification of bouton numbers of MN-Ib and -Is boutons in each condition normalized to wild-type values. ****P<0.0001. Absolute values for normalized data and additional statistical details are summarized in Table S2.

