## Supplementary figures and images for "Botulinum neurotoxin accurately separates tonic vs phasic transmission and reveals heterosynaptic plasticity rules in Drosophila"

### Supplemental Figure 1

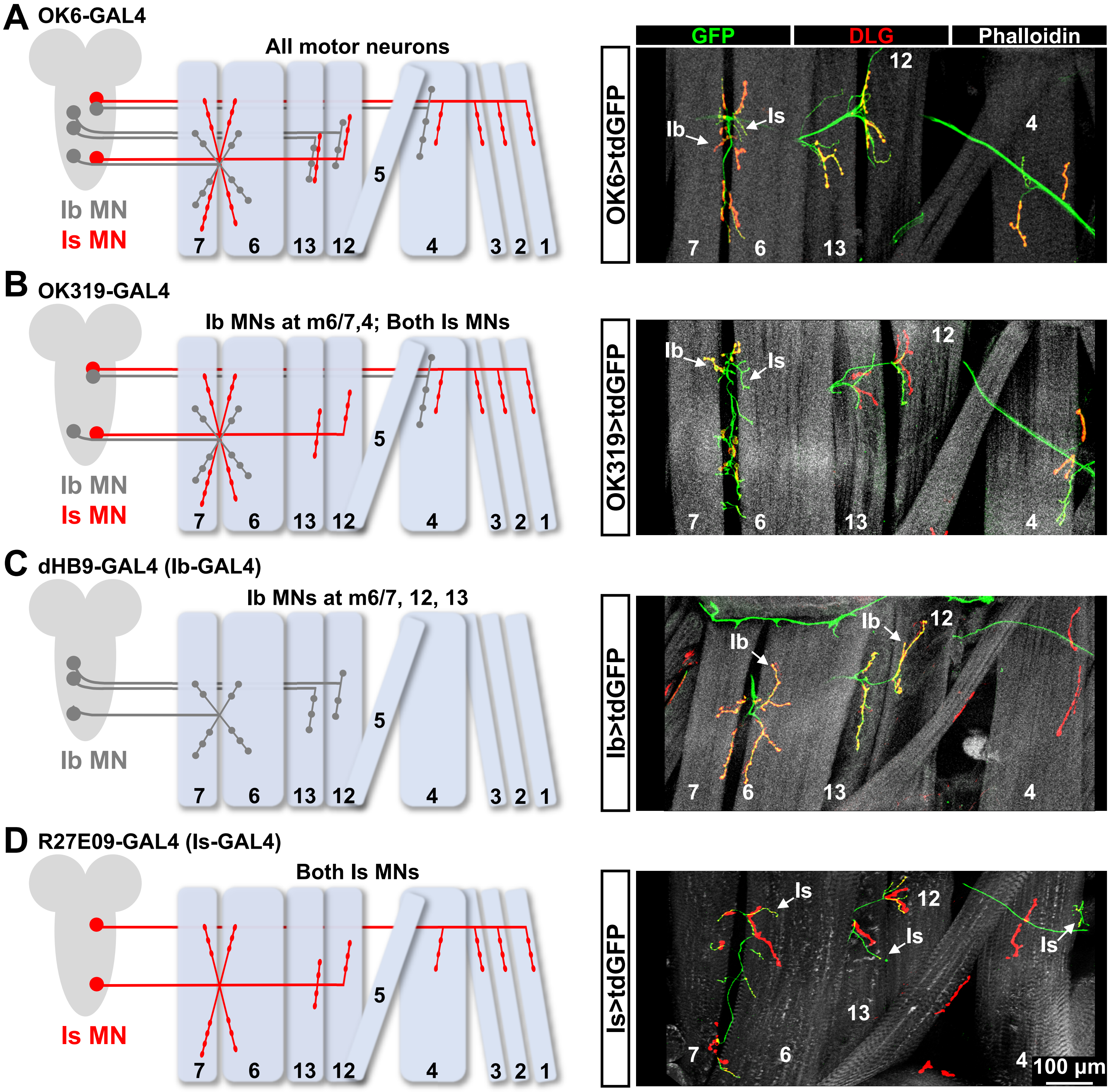


**Figure S1: Motor neuron-specific GAL4 expression at the *Drosophila* NMJ.**

### Supplemental Figure 2

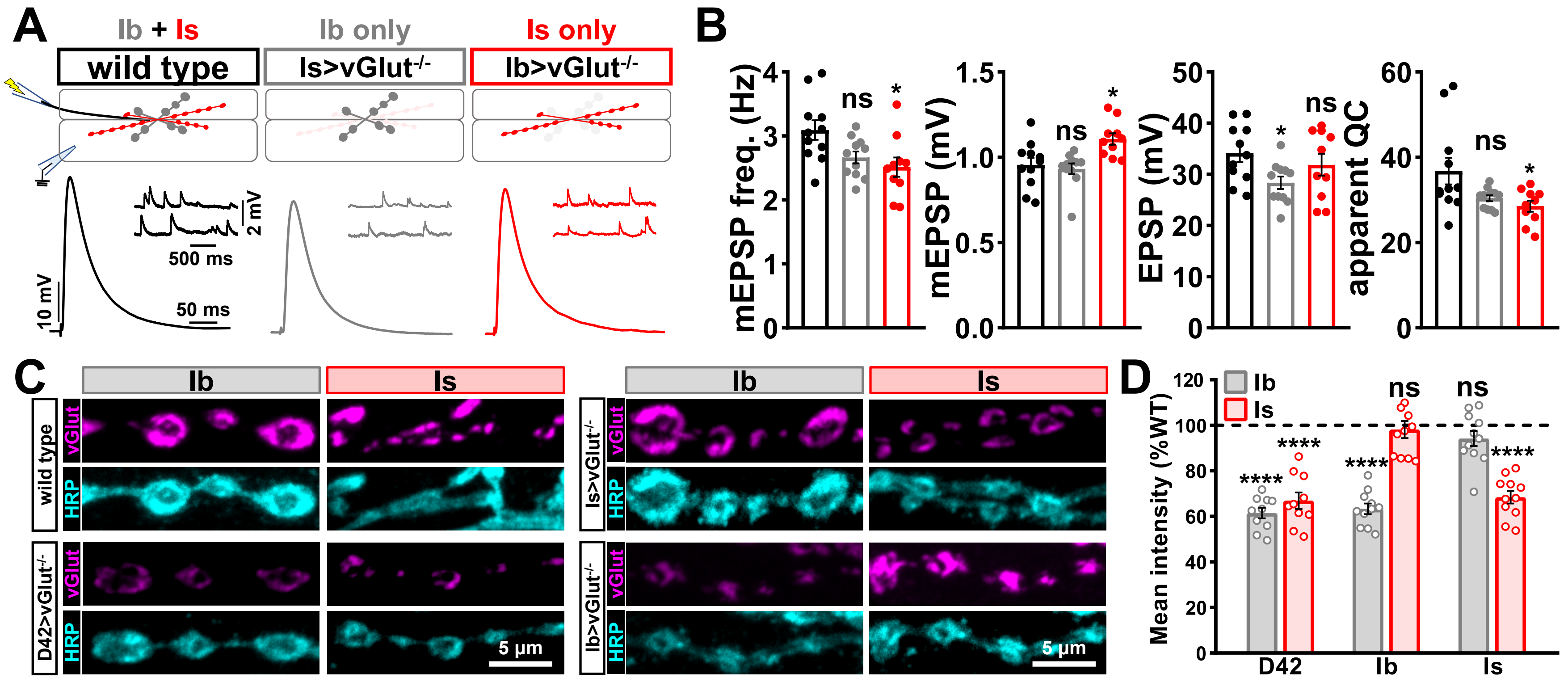


**Figure S2: *vGlut* expression persists despite conditional motor neuron knock out.**

### Supplemental Figure 4

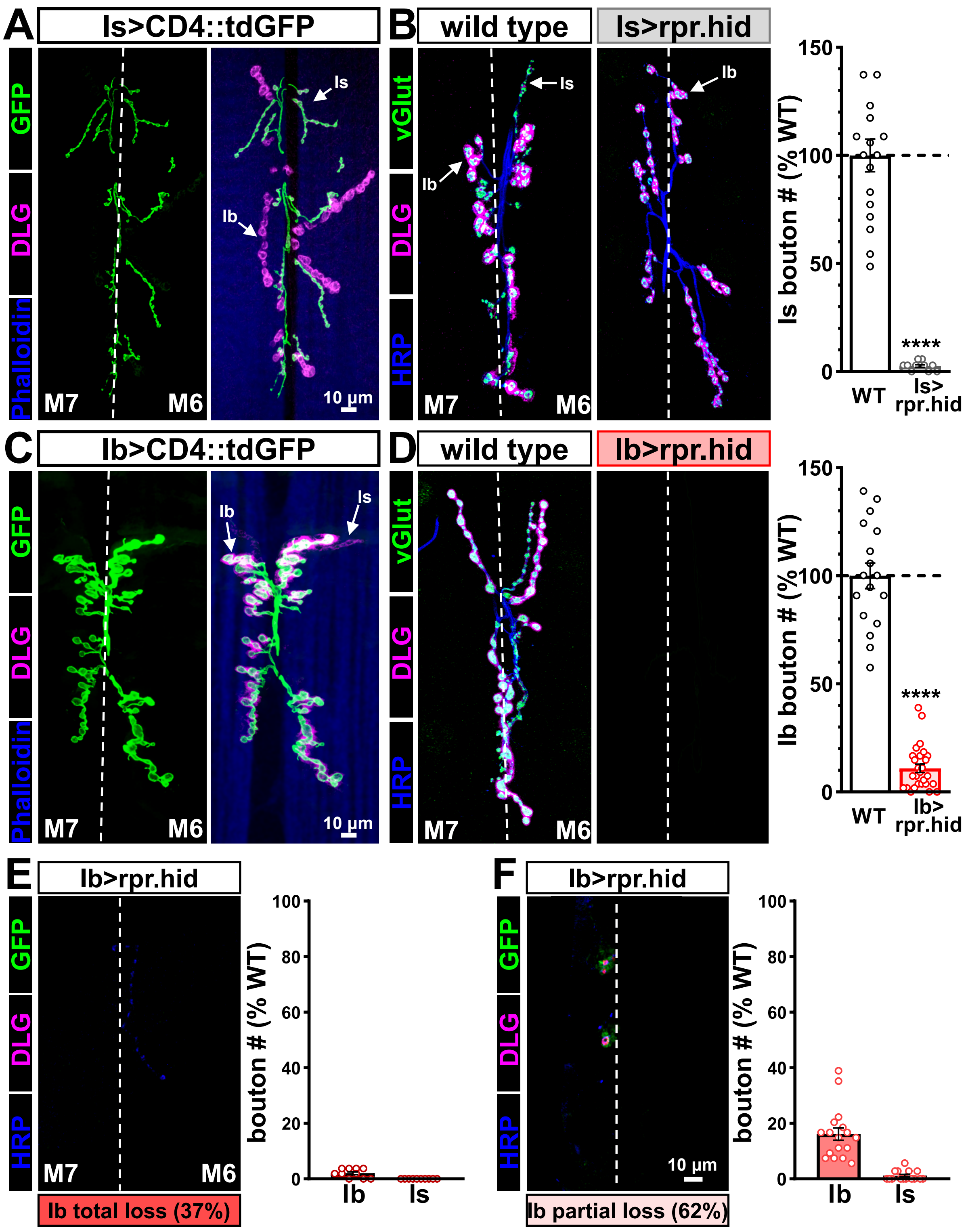


**Figure S4: Genetic ablation of MN-Ib leads to variable ablation of both motor inputs.**
