## Supplemental Figure 3 for "Botulinum neurotoxin accurately separates tonic vs phasic transmission and reveals heterosynaptic plasticity rules in Drosophila"

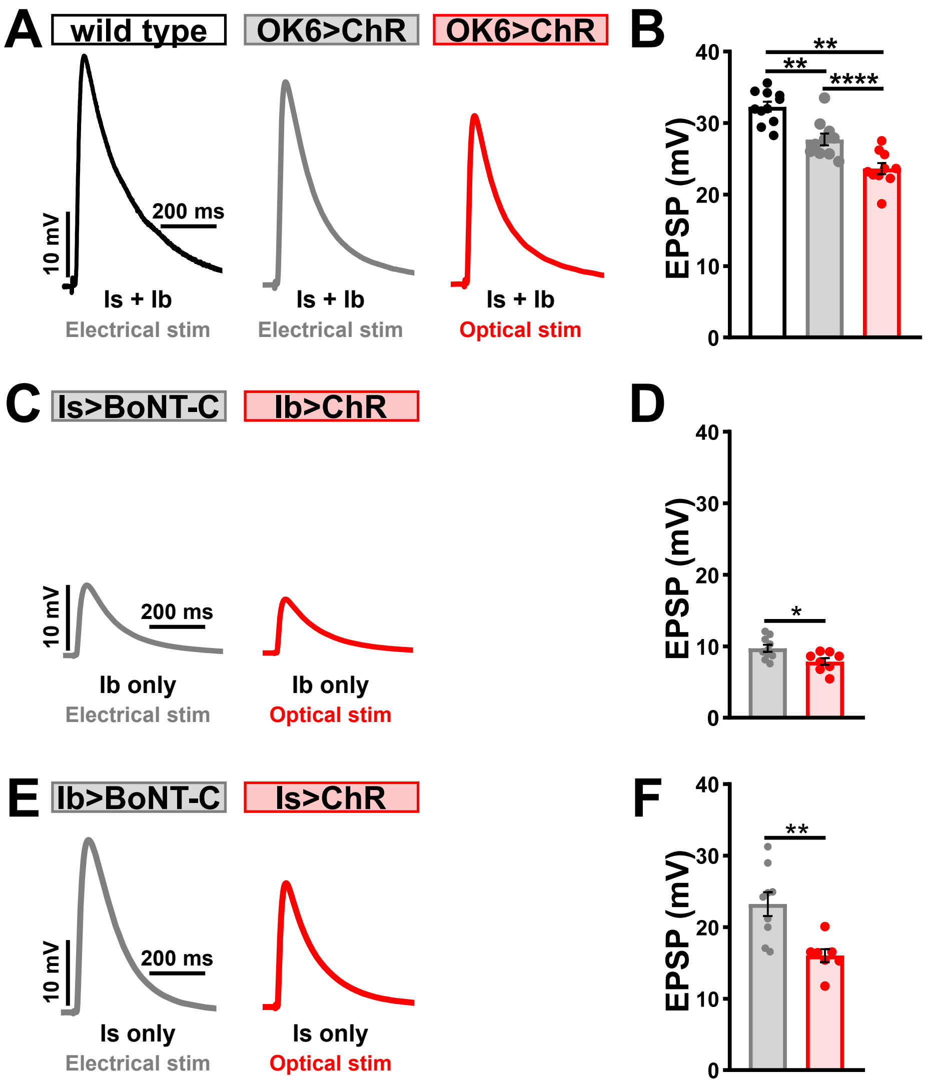


**Figure S3: Electrophysiological differences between optogenetic and electrical stimulation at the *Drosophila* NMJ.**
