## Supplemental Table 1 for "Botulinum neurotoxin accurately separates tonic vs phasic transmission and reveals heterosynaptic plasticity rules in Drosophila"

**Supplementary Table 1: Characterization of synaptic transmission and growth at NMJs expressing BoNT-A, -B, and -E.** All data is related to Figure 2, with genotype, averaged values (with standard error of the mean noted in parentheses), data samples (n), and statistical significance tests and values shown.

| Label | Genotype | mEPSP amplitude (mV) | EPSP amplitude (mV) | QC | mEPSP frequency (Hz) | Rinput (MΩ) | Resting potential (mV) | n | P Value (significance): mEPSP, EPSP, QC, mEPSP freq. |
|---|---|---|---|---|---|---|---|---|---|
| wild type | $w^{1118}$ | 1.01 (±0.037) | 31.46 (±0.841) | 31.61 (±1.166) | 3.25 (±0.128) | 12.01 (±0.136) | 69.14 (±1.115) | 12 | - |
| OK319>BoNT-A | *w*;*OK319-GAL4*/+;*UAS-BoNT-A*/+ | 1.18 (±0.057) | 31.62 (±1.641) | 27.09 (±1.374) | 2.74 (±0.137) | 12.21 (±0.165) | 68.84 (±1.338) | 8 | 0.26 (ns), 0.9999 (ns), 0.26 (ns), 0.08 (ns) |
| OK319>BoNT-B | *w*;*OK319-GAL4*/*UAS-BoNT-B*;+ | 1.48 (±0.129) | 39.34 (±1.858) | 28.01 (±2.561) | 2.27 (±0.203) | 11.81 (±0.278) | 70.47 (±1.454) | 8 | <0.001 (***), <0.01 (**), 0.43 (ns), <0.001 (***) |
| OK319>BoNT-E | *w*;*OK319-GAL4*/+;*UAS-BoNT-E*/+ | 1.06 (±0.069) | 34.28 (±2.554 | 32.94 (±2.671) | 3.35 (±0.177) | 11.98 (±0.143 | 71.03 (±1.306) | 9 | 0.92 (ns), 0.50 (ns), 0.93 (ns), 0.95 (ns) |

| Label | Genotype | Is Bouton #/M6 | Ib Bouton #/M6 | n | P Value (significance): Is Bouton, Ib Bouton |
|---|---|---|---|---|---|
| wild type | $w^{1118}$ | 23.53 (±0.72) | 28.93 (±0.86) | 15 | - |
| OK319>BoNT-A | *w*;*OK319-GAL4*/+;*UAS-BoNT-A*/+ | 24.33 (±1.80) | 28.33 (±2.33) | 12 | 0.97 (ns), 0.99 (ns), |
| OK319>BoNT-B | *w*;*OK319-GAL4*/*UAS-BoNT-B*;+ | 18.17 (±2.66) | 20.58 (±1.25) | 12 | 0.05 (ns), <0.01 (**) |
| OK319>BoNT-E | *w*;*OK319-GAL4*/+;*UAS-BoNT-E*/+ | 22.666 (±1.95) | 27.58 (±2.11) | 12 | 0.96 (ns), 0.97 (ns) |
