## Supplemental Table 2 for "Botulinum neurotoxin accurately separates tonic vs phasic transmission and reveals heterosynaptic plasticity rules in Drosophila"

**Supplementary Table 2: Absolute values for normalized data and additional statistical details.** The figure and panel, genotype, and conditions are noted. Average values (with standard error of the mean noted in parentheses), data samples (n), and statistical significance tests and values are shown for all data.

| Figure | Label | Genotype | mEPSP amplitude (mV) | EPSP amplitude (mV) | QC | mEPSP frequency (Hz) | Rinput (MΩ) | Resting potential (mV) | n | P Value (significance): mEPSP, EPSP, QC, mEPSP freq. |
|---|---|---|---|---|---|---|---|---|---|---|
| 1A,B | OK6>ChR2 | *w;OK6-GAL4/UAS-ChR2$^{T159C}$;+* | 0.98 (±0.04) | 27.72 (±0.82) | 28.76 (±1.73) | 3.28 (±0.057) | 12.11 (±0.24) | -67.97 (±1.20) | 10 | - |
| 1A,B | Ib>ChR2 | *w;UAS-ChR2$^{T159C}$/+;dHb9-GAL4/+* | 1.05 (±0.06) | 11.53 (±0.78) | 11.31 (±1.10) | 3.23 (±0.14) | 11.71 (±0.270) | -68.36 (±0.82) | 8 | <0.0001 (****), 0.611 (ns), <0.0001 (****), 0.956 (ns) |
| 1A,B | Is>ChR2 | *w;UAS-ChR2$^{T159C}$/+;R27E09-GAL4/+* | 1.04 (±0.06) | 16.52 (±0.97) | 16.36 (±1.34) | 3.14 (±0.14) | 12.22 (±0.16) | -68.56 (±1.10) | 9 | <0.0001 (****), 0.722 (ns), <0.0001 (****), 0.709 (ns) |
| 1C,D | wild type | *w$^{1118}$* | 1.05 (±0.06) | 35.70 (±1.75) | 34.99 (±3.10) | 3.21 (±0.12) | 12.23 (±0.24) | -69.84 (±1.49) | 8 | - |
| 1C,D | Is>TNT | *w;UAS-TNT/+;R27E09-GAL4/+* | 0.935 (±0.05) | 8.407 (±0.62) | 8.976 (±0.40) | 3.010 (±0.13) | 12.144 (±0.34) | -70.515 (±1.13) | 9 | <0.0001 (****), 0.258 (ns), <0.0001 (****), 0.488 (ns) |
| 1C,D | Ib>TNT | *w;UAS-TNT/+;dHb9-GAL4/+* | 0.94 (±0.06) | 24.37 (±1.16) | 26.18 (±1.04) | 3.02 (±0.18) | 12.19 (±0.18) | -67.26 (±0.95) | 8 | <0.0001 (****), 0.301 (ns), <0.01 (**), 0.568 (ns) |
| 1E,F | wild type | *w$^{1118}$* | 1.00 (±0.06) | 34.36 (±1.57) | 34.78 (±2.45) | 3.28 (±0.07) | 11.92 (±0.17) | -67.32 (±1.66) | 8 | - |
| 1E,F | Is>rpr.hid | *UAS-rpr.hid/w;+;R27E09-GAL4/+* | 0.76 (±0.01) | 9.16 (±0.69) | 11.99 (±0.86) | 1.59 (±0.19) | 11.93 (±0.18) | -71.93 (±1.53) | 9 | <0.0001 (****), 0.646 (ns), <0.0001 (****), <0.0001 (****) |
| 1E,F | Ib>rpr.hid | *UAS-rpr.hid/w;+;dHb9-GAL4/+* | 0.17 (±0.03) | 0.85 (±0.18) | 2.52 (±0.57) | 0.31 (±0.08) | 12.01 (±0.18) | -70.63 (±0.47) | 10 | <0.0001 (****), 0.607 (ns), <0.0001 (****), <0.0001 (****) |
| 2D,E | wild type | *w$^{1118}$* | 1.04 (±0.08) | 31.78 (±1.14) | 32.46 (±3.38) | 3.267 (±0.11) | 11.72 (±0.13) | -67.16 (±1.95) | 8 | - |
| 2D,E | OK319>TNT | *w;OK319-GAL4/UAS-TNT;+* | 0.94 (±0.04) | 2.25 (±0.43) | 33.97 (±1.93) | 3.13 (±0.11) | 12.17 (±0.16) | -69.91 (±1.28) | 8 | 0.875 (ns), <0.0001 (****), <0.0001 (****), 0.928 (ns) |
| 2D,E | OK319>BoNT-C | *w;OK319-GAL4/+;UAS-BoNT-C/+* | 0.06 (±0.02) | 0.50 (±0.14) | 2.00 (±1.09) | 0.02 (±0.01) | 12.14 (±0.17) | -68.06 (±0.73) | 10 | <0.0001 (****), <0.0001 (****), <0.0001 (****), <0.0001 (****) |
| 7A-E | wild type (seed) | *w$^{1118}$* | 0.99 (±0.05) | 32.6 (±2.19) | 34.19 (±3.74) | 3.26 (±0.15) | 12.20 (±0.18) | -70.00 (±1.06) | 10 | - |
| 7A-E | Is>BoNT-C (seed) | *w;+;R27E09-GAL4/UAS-BoNT-C* | 0.77 (±0.02) | 9.50 (±0.30) | 12.51 (±0.53) | 2.12 (±0.05) | 12.11 (±0.16) | -70.53 (±0.61) | 17 | - |
| 7A-E | Ib>BoNT-C (seed) | *w;+;dHb9-GAL4/UAS-BoNT-C* | 1.34 (±0.04) | 23.65 (±0.48) | 17.92 (±0.62) | 1.51 (±0.05) | 12.05 (±0.14) | -68.60 (±0.65) | 17 | - |
| 7A-E | wild type (reconstituted) | *w$^{1118}$* | 1.00 (±0.001) | 33.10 (±0.026) | 32.70 (±0.043) | 3.50 (±0.003) | - | - | $10^3$ | - |
| 7A-E | Is>BoNT-C (reconstituted) | *w;+;R27E09-GAL4/UAS-BoNT-C* | 0.80 (±0.001) | 9.50 (±0.01) | 12.40 (±0.02) | 2.10 (±0.001) | - | - | $10^3$ | - |
| 7A-E | Ib>BoNT-C (reconstituted) | *w;+;dHb9-GAL4/UAS-BoNT-C* | 1.30 (±0.001) | 23.60 (±0.02) | 17.70 (±0.02) | 1.50 (±0.002) | - | - | $10^3$ | - |
| 7A-E | Ib+Is (reconstituted) | - | 1.00 (±0.001) | 33.10 (±0.02) | 30.10 (±0.02) | 3.50 (±0.002) | - | - | $10^3$ | 0.123 (ns), 0.744 (ns), <0.0001 (****) 0.529 (ns) |
| 8A,B | Is>BoNT-C | *w;+;R27E09-GAL4/UAS-BoNT-C* | 0.78 (±0.02) | 9.13 (±0.78) | 11.59 (±0.67) | 2.12 (±0.12) | 12.02 (±0.18) | -67.54 (±1.59) | 10 | - |
| 8A,B | Is>TNT | *w;UAS-TNT/+;R27E09-GAL4/+* | 0.94 (±0.034) | 8.07 (±0.44) | 8.58 (±0.40) | 2.94 (±0.14) | 12.18 (±0.28) | -69.37 (±1.33) | 9 | <0.05 (*), 0.487 (ns), <0.05 (*), <0.0001 (****) |
| 8A,B | Is>rpr.hid | *UAS-rpr.hid/w;+;R27E09-GAL4/+* | 0.77 (±0.05) | 14.23 (±0.73) | 18.92 (±0.83) | 2.2 (±0.06) | 11.93 (±0.16) | -68.44 (±1.12) | 12 | 0.988 (ns), <0.0001 (****), <0.0001 (****), 0.782 (ns) |
| 8A,B | Ib>BoNT-C | *w;+;dHb9-GAL4/UAS-BoNT-C* | 1.28 (±0.08) | 23.53 (±1.43) | 18.47 (±0.64) | 1.42 (±0.16) | 11.84 (±0.16) | -66.42 (±1.06) | 9 | - |
| 8A,B | Ib>TNT | *w;UAS-TNT/+;dHb9-GAL4/+* | 0.94 (±0.06) | 24.37 (±1.16) | 26.18 (±1.04) | 3.02 (±0.18) | 11.69 (±0.26) | -67.58 (±1.71) | 8 | <0.01 (**), 0.801 (ns), <0.0001 (****), <0.0001 (****) |

| | | | | | | | | | | |
|---|---|---|---|---|---|---|---|---|---|---|
| 8A,B | Ib>rpr.hid | *UAS-rpr.hid/w;+;dHb9-GAL4/+* | 0.19 (±0.05) | 0.93 (±0.17) | 2.16 (±0.66) | 0.26 (±0.06) | 11.98 (±0.14) | -68.50 (±1.03) | 10 | <0.0001 (****), <0.0001 (****), <0.0001 (****), <0.0001 (****) |
| S2B | wild type | $w^{1118}$ | 0.96 (±0.04) | 34.12 (±1.70) | 36.77 (±3.16) | 3.09 (±0.15) | 12.33 (±0.17) | -69.77 (±1.04) | 11 | - |
| S2B | Is>vGlut$^{-/-}$ | *w;UAS-TNT/+;R27E09-GAL4/+* | 0.93 (±0.03) | 28.36 (±1.23) | 30.40 (±0.73) | 2.66 (±0.09) | 12.28 (±0.24) | 71.65 (±0.75) | 11 | <0.0001 (****), 0.258 (ns), <0.0001 (****), 0.488 (ns) |
| S2B | Ib>vGlut$^{-/-}$ | *w;UAS-TNT/+;dHb9-GAL4/+* | 1.11 (±0.03) | 31.88 (±2.12) | 28.56 (±1.30) | 2.51 (±0.15) | 11.52 (±0.23) | -68.94 (±1.12) | 8 | <0.0001 (****), 0.301 (ns), <0.01 (**), 0.568 (ns) |
| S3B | wild type (electrical) | $w^{1118}$ | 1.092 (±0.09) | 32.29 (±0.69) | 28.28 (±2.31) | 3.09 (±0.15) | 11.72 (±0.21) | -68.55 (±1.45) | 11 | - |
| S3B | OK6>ChR2 (electrical) | *w;OK6-GAL4/UAS-ChR2$^{T159C}$;+* | 0.98 (±0.04) | 27.72 (±0.82) | 28.76 (±1.73) | 2.68 (±0.16) | 12.03 (±0.17) | 69.93 (±1.16) | 10 | <0.0001 (****), 0.258 (ns), <0.0001 (****), 0.488 (ns) |
| S3B | OK6>ChR2 (optical) | *w;OK6-GAL4/UAS-ChR2$^{T159C}$;+* | 0.98 (±0.04) | 23.63 (±0.77) | 24.41 (±1.23) | 2.68 (±0.16) | 12.03 (±0.17) | 69.93 (±1.16) | 10 | <0.0001 (****), 0.301 (ns), <0.01 (**), 0.568 (ns) |
| S3D | Is>BoNT-C (electrical) | *w;+;R27E09-GAL4/UAS-BoNT-C* | 0.76 (±0.05) | 9.71 (±0.48) | 13.22 (±0.93) | 2.65 (±0.12) | 11.68 (±0.18) | 68.43 (±0.97) | 10 | <0.0001 (****), 0.258 (ns), <0.0001 (****), 0.488 (ns) |
| S3D | Ib>ChR2 (optical) | *w;UAS-ChR2$^{T159C}$/+;dHb9-GAL4/+* | 1.046 (±0.06) | 7.87 (±0.47) | 7.642 (±0.54) | 2.86 (±0.18) | 11.92 (±0.23) | -70.68 (±0.91) | 8 | <0.0001 (****), 0.301 (ns), <0.01 (**), 0.568 (ns) |
| S3F | Ib>BoNT-C (electrical) | *w;+;dHb9-GAL4/UAS-BoNT-C* | 1.242 (±0.11) | 23.23 (±1.67) | 19.99 (±2.43) | 2.06 (±0.09) | 11.74 (±0.22) | 68.38 (±1.75) | 9 | <0.0001 (****), 0.258 (ns), <0.0001 (****), 0.488 (ns) |
| S3F | Is>ChR2 (optical) | *w;UAS-ChR2$^{T159C}$/+;R27E09-GAL4/+* | 1.04 (±0.06) | 16.15 (±0.74) | 16.00 (±1.17) | 2.74 (±0.17) | 12.08 (±0.16) | -68.46 (±1.16) | 9 | <0.0001 (****), 0.301 (ns), <0.01 (**), 0.568 (ns) |

| Figure | Label | Genotypes | NMJ | Bouton #/M6 | n | BRP puncta #/M6 | BRP puncta intensity/M6 (%WT) | n | P Value (significance): Bouton#, BRP#, BRP intensity |
|---|---|---|---|---|---|---|---|---|---|
| 3A,B | wild type | $w^{1118}$ | Ib | 28.50 (±1.21) | 16 | 217.33 (±9.61) | 100 (±4.93) | 12 | - |
| 3A,B | OK319>TNT | *w;OK319-GAL4/UAS-TNT;+* | Ib | 23.09 (±0.77) | 11 | 192.58 (±4.56) | 98.13 (±4.41) | 12 | <0.05 (*), <0.05 (*), 0.933 (ns) |
| 3A,B | OK319>BoNT-C | *w;OK319-GAL4/+;UAS-BoNT-C/+* | Ib | 27.92 (±2.14) | 12 | 210.58 (±5.71) | 97.79 (±3.45) | 12 | 0.944 (ns), 0.719 (ns), 0.908 (ns) |
| 3A,B | wild type | $w^{1118}$ | Is | 23.19 (±0.56) | 16 | 93.58 (±3.70) | 100 (±3.93) | 12 | - |
| 3A,B | OK319>TNT | *w;OK319-GAL4/UAS-TNT;+* | Is | 28.55 (±0.95) | 11 | 111.17 (±6.24) | 102.97 (±4.75) | 12 | <0.05 (*), <0.05 (*), 0.810 (ns) |
| 3A,B | OK319>BoNT-C | *w;OK319-GAL4/+;UAS-BoNT-C/+* | Is | 23.58 (±2.23) | 12 | 99.50 (±3.86) | 93.06 (±2.63) | 12 | 0.968 (ns), 0.586 (ns), 0.352 (ns) |

| Figure | Label | Genotypes | T-bar length (nm) | Active zone length (nm) | Vesicle density (#/µm$^2$) | n | P Value (significance): T-bar length, AZ length, Vesicle density |
|---|---|---|---|---|---|---|---|
| 3C,D | wild type | $w^{1118}$ | 122.29 (±2.85) | 584.78 (±14.48) | 171.32 (±5.69) | 18 | - |
| 3C,D | OK319>TNT | *w;OK319-GAL4/UAS-TNT;+* | 122.76 (±3.61) | 522.93 (±14.94) | 183.51 (±4.48) | 20 | <0.05 (*), <0.05 (*), 0.195 (ns) |
| 3C,D | OK319>BoNT-C | *w;OK319-GAL4/+;UAS-BoNT-C/+* | 123.07 (±3.64) | 555.47 (±17.40) | 168.51 (±5.76) | 19 | 0.944 (ns), 0.719 (ns), 0.904 (ns) |

| Figure | Label | Genotypes | NMJ | GluRIIA puncta intensity (% wild type) | GluRIIB puncta intensity (% wild type) | GluRIID puncta intensity (% wild type) | n | P Value (significance): GluRIIA, GluRIIB, GluRIID |
|---|---|---|---|---|---|---|---|---|
| 4B | wild type | *$w^{1118}$* | Ib | 100 (±3.70) | 100 (±4.09) | 100 (±2.56) | 16 | - |
| 4B | wild type | *$w^{1118}$* | Is | 67.28 (±2.78) | 121.53 (±7.87) | 94.26 (±3.84) | 15 | <0.0001 (****), <0.05 (*), 0.218 (ns) |
| 4E | OK319>BoNT-C | *w;OK319-GAL4/+;UAS-BoNT-C/+* | Ib | 100 (±3.12) | 100 (±2.60) | 100 (±4.38) | 14 | - |
| 4E | OK319>BoNT-C | *w;OK319-GAL4/+;UAS-BoNT-C/+* | Is | 68.25 (±3.88) | 83.73 (±2.18) | 93.89 (±4.31) | 14 | <0.0001 (****), <0.01 (**), 0.329 (ns) |

| Figure | Label | Genotypes | NMJ | Transmission | Quantal size (ΔF/F) | n | P Value (significance): |
|---|---|---|---|---|---|---|---|
| 5B | Is>BoNT-C | *w;MHC>GCaMP8f/+;R27E09-GAL4/UAS-BoNT-C* | Ib | spontaneous | 0.029 (0.002) | 5 | - |
| 5B | Is>BoNT-C | *w;MHC>GCaMP8f/+;R27E09-GAL4/UAS-BoNT-C* | Is | spontaneous | 0.004 (0.001) | 5 | <0.0001 (****) |
| 5C | Is>BoNT-C | *w;MHC>GCaMP8f/+;R27E09-GAL4/UAS-BoNT-C* | Ib | evoked | 0.326 (±0.018) | 5 | - |
| 5C | Is>BoNT-C | *w;MHC>GCaMP8f/+;R27E09-GAL4/UAS-BoNT-C* | Is | evoked | 0.013 (±0.001) | 5 | <0.0001 (****) |
| 5E | Ib>BoNT-C | *w;MHC>GCaMP8f/+;dHb9-GAL4/UAS-BoNT-C* | Ib | spontaneous | 0.005 (±0.001) | 5 | - |
| 5E | Ib>BoNT-C | *w; MHC>GCaMP8f/+;dHb9-GAL4/UAS-BoNT-C* | Is | spontaneous | 0.046 (±0.004) | 5 | <0.0001 (****) |
| 5F | Ib>BoNT-C | *w;MHC>GCaMP8f/+;dHb9-GAL4/UAS-BoNT-C* | Ib | evoked | 0.013 (±0.001) | 5 | - |
| 5F | Ib>BoNT-C | *w; MHC>GCaMP8f/+;dHb9-GAL4/UAS-BoNT-C* | Is | evoked | 0.475 (±0.026) | 5 | <0.0001 (****) |

| Figure | Label | Genotype | Muscle | Is Bouton #/NMJ | Ib Bouton #/NMJ | n | P Value (significance): Is Bouton, Ib Bouton |
|---|---|---|---|---|---|---|---|
| 6A-D | wild type | *$w^{1118}$* | 6 | 23.53 (±0.72) | 28.33 (±0.86) | 15 | - |
| 6A-D | Is>BoNT-C | *w;+;R27E09-GAL4/UAS-BoNT-C* | 6 | 24.33 (±1.80) | 25.56 (±0.79) | 16 | 0.993 (ns), 0.999 (ns), |
| 6A-D | Is>TNT | *w;UAS-TNT/+;R27E09-GAL4/+* | 6 | 27.83 (±1.65) | 27.67 (±1.52) | 12 | <0.05 (*), 0.811 (ns) |
| 6A-D | Is>rpr.hid | *UAS-rpr.hid/w;+;R27E09-GAL4/+* | 6 | 31.73 (±0.83) | 31.73 (±0.83) | 11 | <0.0001 (****), <0.001 (***) |
| 6E-H | wild type | *$w^{1118}$* | 6 | 23.23 (±0.48) | 27.15 (±0.68) | 13 | - |
| 6E-H | Ib>BoNT-C | *w;+;dHb9-GAL4/UAS-BoNT-C* | 6 | 23.69 (±0.71) | 25.63 (±0.65) | 16 | 0.999 (ns), 0.999 (ns), |
| 6E-H | Ib>TNT | *w;UAS-TNT/+;dHb9-GAL4/+* | 6 | 22.09 (±1.56) | 21.08 (±0.81) | 11 | 0.982 (ns), <0.01 (**) |
| 6E-H | Ib>rpr.hid | *UAS-rpr.hid/w;+;dHb9-GAL4/+* | 6 | 0.28 (±0.11) | 5.44 (±0.85) | 25 | <0.0001 (****), <0.0001 (****) |

| Figure | Label | Genotypes | Ib vGlut puncta intensity (% wild type) | Is vGlut puncta intensity (% wild type) | n | P Value (significance): Ib vGlut, Is vGlut |
|---|---|---|---|---|---|---|
| S2D | wild type | *$w^{1118}$* | 100 (±3.59) | 100 (±3.47) | 10 | - |
| S2D | D42>vGlut$^{-/-}$ | *w;vGlut$^{SS1}$/B3RT-vGlut-B3RT;UAS-B3/D42-GAL4* | 61.49 (±2.33) | 61.49 (±3.60) | 10 | <0.0001 (****), <0.0001 (****) |
| S2D | Is>vGlut$^{-/-}$ | *w;vGlut$^{SS1}$/B3RT-vGlut-B3RT;UAS-B3/R7E09-GAL4* | 94.24 (±3.36) | 68.46 (±2.71) | 11 | 0.384 (ns), <0.0001 (****) |
| S2D | Ib>vGlut$^{-/-}$ | *w;vGlut$^{SS1}$/B3RT-vGlut-B3RT;UAS-B3/dHB9-GAL4* | 63.36 (±2.33) | 98.21 (±3.77) | 11 | <0.0001 (****), 0.965 (ns) |

| Figure | Label | Genotype | Muscle 6/7 Is Bouton #/NMJ | Muscle 6/7 Ib Bouton #/NMJ | n | P Value (significance): Is Bouton, Ib Bouton |
|---|---|---|---|---|---|---|
| S4A,B | wild type | *w*[1118] | 34.94 (±4.07) | 53.88 (±3.19) | 17 | - |
| S4A,B | Is>rpr.hid | *UAS-rpr.hid/w;+;R27E09-GAL4/+* | 0.91 (±0.21) | 61.91 (±2.51) | 11 | <0.0001 (****), <0.05 (*) |
| S4A,B | Ib>rpr.hid | *UAS-rpr.hid/w;+;dHb9-GAL4/+* | 0.22 (±0.09) | 5.89 (±1.05) | 27 | <0.0001 (****), <0.0001 (****) |
| S4E,F | Ib>rpr.hid (total loss) | *UAS-rpr.hid/w;+;dHb9-GAL4/+* | 0 (±0) | 1.10 (±0.28) | 10 (37%) | - |
| S4E,F | Ib>rpr.hid (partial loss) | *UAS-rpr.hid/w;+;dHb9-GAL4/+* | 0.35 (±0.15) | 8.71 (±1.22) | 17 (63%) | - |

| Figure | Label | Genotype | Muscle | mEPSP amplitude (mV) | EPSP amplitude (mV) | QC | mEPSP frequency (Hz) | Rinput (MΩ) | Resing potential (mV) | n | P Value (significance): mEPSP, EPSP, QC, mEPSP freq. |
|---|---|---|---|---|---|---|---|---|---|---|---|
| | wild type | *w*[1118] | 6 | 1.00 (±0.05) | 32.60 (±2.19) | 34.19 (±3.74) | 3.26 (±0.15) | 12.48 (±0.22) | -68.80 (±1.78) | 10 | - |
| | Is>BoNT-C | *w;+;R27E09-GAL4/UAS-BoNT-C* | 6 | 0.78 (±0.02) | 9.13 (±0.78) | 11.59 (±0.67) | 2.12 (±0.12) | 11.61 (±0.14) | -70.62 (±0.65) | 10 | 0.261 (ns), 0.999 (ns), 0.256 (ns), 0.077 (ns) |
| | Is>TNT | *w;UAS-TNT/+;R27E09-GAL4/+* | 6 | 0.94 (±0.03) | 8.07 (±0.44) | 8.59 (±0.40) | 2.94 (±0.14) | 11.95 (±0.15) | -69.43 (±1.27) | 9 | <0.001 (***), <0.01 (**), 0.434 (ns), <0.001 (***) |
| | Is>rpr.hid | *UAS-rpr.hid/w;+;R27E09-GAL4/+* | 6 | 0.77 (±0.05) | 14.23 (±0.73) | 18.92 (±0.83) | 2.2 (±0.055) | 11.86 (±0.11) | -68.85 (±0.88) | 12 | 0.919 (ns), 0.499 (ns), 0.928 (ns), 0.949 (ns) |
| | Ib>BoNT-C | *w;+;exex-GAL4/UAS-BoNT-C* | 6 | 1.28 (±0.08) | 23.53 (±1.43) | 18.47 (±0.64) | 1.42 (±0.16) | 11.77 (±0.17) | -65.65 (±1.35) | 9 | 0.2613 (ns), 0.999 (ns), 0.256 (ns), 0.076 (ns) |
| | Ib>TNT | *w;UAS-TNT/+;exex-GAL4/+* | 6 | 0.94 (±0.06) | 24.37 (±1.16) | 26.18 (±1.04) | 3.02 (±0.18) | 12.10 (±0.19) | -68.40 (±1.308) | 8 | <0.001 (***), <0.01 (**), 0.434 (ns), <0.001 (***) |
| | Ib>rpr.hid | *UAS-rpr.hid/w;+;exex-GAL4/+* | 6 | 0.19 (±0.05) | 0.93 (±0.17) | 2.16 (±0.66) | 0.26 (±0.06) | 11.81 (±0.16) | -69.71 (±1.05) | 10 | 0.919 (ns), 0.499 (ns), 0.928 (ns), 0.949 (ns) |
| | wild type | *w*[1118] | 12 | 0.98 (±0.06) | 30.92 (±1.41) | 31.86 (±0.82) | 3.16 (±0.19) | 12.02 (±0.27) | -69.72 (±0.78) | 12 | - |
| | Is>BoNT-C | *w;+;R27E09-GAL4/UAS-BoNT-C* | 12 | 0.78 (±0.04) | 9.82 (±0.50) | 12.73 (±0.34) | 2.14 (±0.21) | 12.07 (±0.15) | -70.01 (±0.79) | 8 | <0.05 (*), <0.0001 (****), <0.0001 (****), <0.001 (***) |
| | Is>TNT | *w;UAS-TNT/+;R27E09-GAL4/+* | 12 | 0.95 (±0.06) | 10.50 (±0.76) | 11.13 (±0.39) | 2.76 (±0.14) | 12.3 (±0.244) | -69.02 (±1.21) | 8 | 0.957 (ns), <0.0001 (****), <0.0001 (****), 0.377 (ns) |
| | Is>rpr.hid | *UAS-rpr.hid/w;+;R27E09-GAL4/+* | 12 | 0.76 (±0.03) | 12.66 (±0.62) | 16.72 (±0.52) | 2.08 (±0.12) | 12.17 (±0.15) | -69.29 (±1.45) | 9 | <0.05 (*), <0.0001 (****), <0.0001 (****), <0.001 (***) |
| | wild type | *w*[1118] | 4 | 1.01 (±0.05) | 27.69 (±1.35) | 28.25 (±2.08) | 3.13 (±0.09) | 12.05 (±0.11) | -69.48 (±0.94) | 11 | - |
| | Is>BoNT-C | *w;+;R27E09-GAL4/UAS-BoNT-C* | 4 | 0.75 (±0.03) | 13.25 (±0.66) | 17.95 (±1.11) | 2.19 (±0.08) | 12.32 (±0.21) | -70.02 (±0.96) | 12 | <0.001 (***), <0.0001 (****), <0.0001 (****), <0.0001 (****) |
| | Is>TNT | *w;UAS-TNT/+;R27E09-GAL4/+* | 4 | 1.04 (±0.05) | 13.55 (±0.64) | 13.41 (±1.09) | 2.84 (±0.15) | 11.91 (±0.18) | -67.15 (±0.79) | 12 | 0.949 (ns), <0.0001 (****), <0.0001 (****), 0.2210 (ns) |
| | Is>rpr.hid | *UAS-rpr.hid/w;+;R27E09-GAL4/+* | 4 | 0.79 (±0.03) | 16.24 (±0.99) | 20.89 (±1.28) | 2.07 (±0.08) | 11.68 (±0.19) | -69.42 (±0.83) | 12 | <0.01 (**), <0.0001 (****), <0.01 (**), <0.0001 (****) |

| Figure | Label | Genotype | Muscle | Is Bouton #/NMJ | Significant Change (%WT) | Ib Bouton #/NMJ | Significant Change (%WT) | n | P Value (significance): Is Bouton, Ib Bouton |
|---|---|---|---|---|---|---|---|---|---|
| | wild type | $w^{1118}$ | 6 | 23.53 (±0.72) | - | 28.33 (±0.86) | - | 15 | - |
| | Is>BoNT-C | *w;+;R27E09-GAL4/UAS-BoNT-C* | 6 | 24.33 (±1.80) | - | 25.56 (±0.79) | - | 16 | 0.993 (ns), 0.999 (ns), |
| | Is>TNT | *w;UAS-TNT/+;R27E09-GAL4/+* | 6 | 27.83 (±1.65) | ↑24 | 27.67 (±1.52) | - | 12 | <0.05 (*), 0.811 (ns) |
| | Is>rpr.hid | *UAS-rpr.hid/w;+;R27E09-GAL4/+* | 6 | 31.73 (±0.83) | ↓100 | 31.73 (±0.83) | ↑18 | 11 | <0.0001 (****), <0.001 (***) |
| | Ib>BoNT-C | *w;+;exex-GAL4/UAS-BoNT-C* | 6 | 23.69 (±0.71) | - | 25.63 (±0.65) | - | 16 | 0.999 (ns), 0.999 (ns), |
| | Ib>TNT | *w;UAS-TNT/+;exex-GAL4/+* | 6 | 22.09 (±1.56) | - | 21.08 (±0.81) | ↓22 | 11 | 0.982 (ns), <0.01 (**) |
| | Ib>rpr.hid | *UAS-rpr.hid/w;+;exex-GAL4/+* | 6 | 0.28 (±0.11) | ↓99 | 5.44 (±0.85) | ↓94 | 25 | <0.0001 (****), <0.0001 (****) |
| | wild type | $w^{1118}$ | 7 | 12.63 (±1.29) | - | 18.81 (±1.41) | - | 15 | - |
| | Is>BoNT-C | *w;+;R27E09-GAL4/UAS-BoNT-C* | 7 | 12.13 (±0.53) | - | 19.88 (±0.58) | - | 16 | 0.879 (ns), 0.6678 (ns), |
| | Is>TNT | *w;UAS-TNT/+;R27E09-GAL4/+* | 7 | 18.92 (±1.51) | ↑49 | 18.58 (±2.10) | - | 12 | <0.01 (**), 0.993 (ns) |
| | Is>rpr.hid | *UAS-rpr.hid/w;+;R27E09-GAL4/+* | 7 | 0 (±0) | ↓100 | 22.64 (±1.73) | ↑20 | 11 | <0.0001 (****), 0.991 (ns) |
| | Ib>BoNT-C | *w;+;exex-GAL4/UAS-BoNT-C* | 7 | 12.13 (±0.41) | - | 19.38 (±0.78) | - | 16 | 0.879 (ns), 0.889 (ns), |
| | Ib>TNT | *w;UAS-TNT/+;exex-GAL4/+* | 7 | 13.73 (±1.51) | - | 12.82 (±1.23) | ↓30 | 11 | 0.814 (ns), <0.05 (*) |
| | Ib>rpr.hid | *UAS-rpr.hid/w;+;exex-GAL4/+* | 7 | 0 (±0) | ↓99 | 0.44 (±0.18) | ↓99 | 25 | <0.0001 (****), <0.0001 (****) |
| | wild type | $w^{1118}$ | 12 | 23.75 (±0.993) | - | 18.08 (±0.87) | - | 12 | - |
| | Is>BoNT-C | *w;+;R27E09-GAL4/UAS-BoNT-C* | 12 | 24.64 (±2.76) | - | 17.82 (±2.03) | - | 11 | 0.993 (ns), 0.999 (ns), |
| | Is>TNT | *w;UAS-TNT/+;R27E09-GAL4/+* | 12 | 18.92 (±1.51) | ? | 18.58 (±2.10) | - | 12 | <0.05 (*), 0.999 (ns) |
| | Is>rpr.hid | *UAS-rpr.hid/w;+;R27E09-GAL4/+* | 12 | 0 (±0) | ↓100 | 26.82 (±1.48) | ↑48 | 11 | <0.0001 (****), <0.001 (***) |
| | wild type | $w^{1118}$ | 13 | 15.50 (±0.48) | - | 15.33 (±0.41) | - | 12 | - |
| | Is>BoNT-C | *w;+;R27E09-GAL4/UAS-BoNT-C* | 13 | 15.55 (±0.62) | - | 15.91 (±0.41) | - | 11 | 0.999 (ns), 0.793 (ns), |
| | Is>TNT | *w;UAS-TNT/+;R27E09-GAL4/+* | 13 | 21.83 (±0.81) | ↑40 | 14.83 (±0.53) | - | 12 | <0.0001 (****), 0.843 (ns) |
| | Is>rpr.hid | *UAS-rpr.hid/w;+;R27E09-GAL4/+* | 13 | 0 (±0) | ↓100 | 20.73 (±0.74) | ↑35 | 11 | <0.0001 (****), <0.0001 (****) |
| | wild type | $w^{1118}$ | 4 | 22.58 (±0.99) | - | 18.67 (±1.39) | - | 12 | - |
| | Is>BoNT-C | *w;+;R27E09-GAL4/UAS-BoNT-C* | 4 | 24.64 (±2.33) | - | 20.91 (±1.46) | - | 11 | 0.872 (ns), 0.837 (ns), |
| | Is>TNT | *w;UAS-TNT/+;R27E09-GAL4/+* | 4 | 28.83 (±1.17) | ↑27 | 19.67 (±1.66) | - | 12 | <0.05 (*), 0.995 (ns) |
| | Is>rpr.hid | *UAS-rpr.hid/w;+;R27E09-GAL4/+* | 4 | 0 (±0) | ↓100 | 27.27 (±1.24) | ↑45 | 11 | <0.0001 (****), <0.01 (**) |
