## Supplemental Table 3 for "Botulinum neurotoxin accurately separates tonic vs phasic transmission and reveals heterosynaptic plasticity rules in Drosophila"

### STAR★METHODS

#### KEY RESOURCE TABLE

| REAGENT/RESOURCE | | SOURCE | IDENTIFIER |
|---|---|---|---|
| **Antibodies** | **Dilution** | | |
| Mouse anti-GluRIIA (8B4D2) | 1:50 | Developmental Studies Hybridoma Bank (DSHB) | AB_528269 |
| Mouse anti-BRP (nc82) | 1:100 | DSHB | AB_2314866 |
| Mouse anti-DLG (4F3) | 1:100 | DSHB | AB_528203 |
| Mouse anti-GFP (4C9) | 1:500 | DSHB | AB_2617422 |
| Rabbit anti-DLG | 1:10000 | (Pielage et al., 2005) | |
| Rabbit anti-GluRIIB | 1:1000 | (Perry et al., 2017) | |
| Guinea pig anti-vGlut | 1:2000 | (Goel and Dickman, 2018) | |
| Guinea pig anti-GluRIID | 1:1000 | (Perry et al., 2017) | |
| Alexa Fluor 647 conjugated Goat anti-Horseradish Peroxidase | 1:400 | Jackson ImmunoResearch Laboratories (Jackson) | 123-605-021 |
| Alexa Fluor 488 conjugated secondary antibodies | 1:400 | Jackson | 706-545-148, 715-545-150, 711-545-152 |
| Cy3-conjugated secondary antibodies | 1:400 | Jackson | 706-165-148, 715-165-150, 711-165-152 |
| DyLight 405-conjugated secondary antibodies | 1:400 | Jackson | 706-475-148, 715-475-150 |
| Alexa Fluor 647 conjugated Goat anti-Phalloidin | 1:1000 | ThermoFisher | A22287 |
| **Drosophila Strains** | | | |
| UAS-BoNT-C | | This study | |
| UAS-BoNT-A | | This study | |
| UAS-BoNT-B | | This study | |
| UAS-BoNT-E | | This study | |
| MHC-CD8-GCaMP8f-Sh (SynapGCaMP8f) | | This study | |
| OK6-GAL4 | | (Aberle et al., 2002) | |
| UAS-rpr.hid | | (Zhou et al., 1997) | |
| OK319-GAL4 | | (Sweeney et al., 1995) | |
| vGlut$^{SS1}$ | | (Sherer et al., 2020) | |
| B3RT-vGlut-B3RT | | (Sherer et al., 2020) | |
| UAS-B3 | | (Sherer et al., 2020) | |
| UAS-ChR2$^{T159C}$ | | Bloomington Drosophila Stock Center (BDSC) | 58373 |
| UAS-CD4::tdGFP | | BDSC | 35839 |
| dHb9-GAL4 | | BDSC | 83004 |
| GMR27E09-GAL4 | | BDSC | 49227 |
| UAS-TNT | | BDSC | 28838 |
| $w^{1118}$ | | BDSC | 5905 |
| D42-GAL4 | | BDSC | 8816 |
| **Construct and reagents** | | | |
| pUASt destination vector | | Drosophila Genetics Resource Center (DGRC) | 1129 |
| all trans-Retinal | | Sigma-Aldrich | R2500 |
| **Software and Algorithms** | | | |
| NIS Elements software | | Nikon | 4.51.01 |
| Axon pCLAMP Clampfit | | Molecular Devices | 10.7 |
| MiniAnalysis | | Synaptosoft | 6.0.3 |
| GraphPad Prism | | GraphPad | 8.0.1 |
| Jupyter Notebook | | Anaconda | 6.0.1 |
| ImageJ (Fiji) | | (Rueden et al., 2017) | |
